# Hierarchical recruitment of competition alleviates working memory overload in a fronto-parietal model

**DOI:** 10.1101/181370

**Authors:** Dominic Standage, Martin Paré, Gunnar Blohm

**Author notes:** **Correspondence** Dr. Dominic Standage University of Birmingham Hills Building, Room 207 Birmingham, B15 2TT, UK.

## Abstract

The storage limitations of visual working memory have been the subject of intense research interest for several decades, but few studies have systematically investigated the dependence of these limitations on memory load that exceeds our retention abilities. Under this real-world scenario, performance typically declines beyond a critical load among low-performing subjects, a phenomenon known as working memory overload. We used a fronto-parietal cortical model to test the hypothesis that high-performing subjects select a manageable number of items for storage, thereby avoiding overload. The model accounts for behavioural and electrophysiological data from high-performing subjects in a parameter regime where competitive encoding in its prefrontal network selects items for storage, inter-areal projections sustain their representations after stimulus offset, and weak dynamics in its parietal network limit their mutual interference. Violation of these principles accounts for these data among low-performing subjects, implying that poor visual working memory performance reflects poor control over fronto-parietal circuitry, and making testable predictions for experiments.

## 1 Introduction

The transient retention and manipulation of information is central to cognition and is known as working memory. The modular nature of working memory has long been recognized [see Baddeley (2012)], with visual working memory (WM) receiving considerable attention for several decades. Much of this research has focused on the storage limitations of WM, which have been described in terms of the number of stored items and the precision of reports about them [see Luck and Vogel (2013); Ma, Husain, and Bays (2014)]. A large body of work provides evidence for a severe limit on the number of items humans can store in WM, typically three of four memoranda among healthy young adults [see Vogel and Awh (2008); Luck and Vogel (2013)]. This number is often referred to as WM capacity [though see Brady, Konkle, and Alvarez (2011) for a broader definition of this term] and is a reliable predictor of cognitive ability more generally [see Unsworth, Fukuda, Awh, and Vogel (2014)]. Thus, understanding its neural basis is widely regarded as a fundamental goal of cognitive neuroscience.

In the laboratory, the storage limitations of WM are estimated by varying the number of items for retention over a memory delay (WM load), but little emphasis has been given to the dependence of storage limitations on load that exceeds our retention abilities. This real-world scenario has long been of concern to instructional designers (Sweller, 1988; Merrienboër & Sweller, 2005), who consider the avoidance of ‘overload’ a fundamental principle of effective design. Consistent with this concern, recent WM studies have shown a decrease in WM capacity beyond a critical load (Chee & Chuah, 2007; Xu, 2007; Cusack, Lehmann, Veldsman, & Mitchell, 2009; Linke, Vicente-Grabovetsky, Mitchell, & Cusack, 2011; Matsuyoshi, Osaka, & Osaka, 2014; Fukuda, Woodman, & Vogel, 2015), referred to as WM overload (Matsuyoshi et al., 2014). To the best of our knowledge, overload has not been reported in terms of WM precision and we limit our focus to the available data. For a thorough treatment of the mechanistic relationship between WM capacity and precision, see Standage and Paré (2018).

It is widely believed that WM storage is supported by ‘attractor states’ in neocortex, where regenerative excitation sustains neural firing after stimulus offset, kept in check by feedback inhibition [see X.-J. Wang (2001)]. In models of this kind, overload is a consequence of the competition imposed by inhibition (Edin et al., 2009), but not all subjects show overload [*e.g.* Fukuda, Woodman, and Vogel (2015)] and among those who do, overload is not typically as pronounced as in these models [*e.g.* Wei, Wang, and Wang (2012); Section 3.1 here]. The occurrence of overload in attractor models, however, assumes that all items in a stimulus array are encoded for storage. Thus, a viable strategy for managing overload is to limit the number of encoded items (Cusack et al., 2009). We hypothesize that this selection process is implemented by strong competitive dynamics during stimulus encoding.

To test our hypothesis, we simulated a multiple-item WM task with biophysical models of posterior parietal cortex (PPC) and lateral prefrontal cortex (PFC), both of which are extensively correlated with WM [see Curtis (2006); Funahashi (2013)]. PPC is hypothesized to be the hub of distributed WM storage (Palva, Monto, Kulashekhar, & Palva, 2010; Christophel, Hebart, & Haynes, 2012; Salazar, Dotson, Bressler, & Gray, 2012) and is well characterized by neural data from visual tasks [see Goldberg, Bisley, Powell, and Gottlieb (2006); Serences and Yantis (2006)], so we first sought to determine whether competitive encoding in a local-circuit PPC model could alleviate overload in a manner consistent with behavioural data from high-performing WM subjects, and with neural data from visual tasks. We reasoned that any inconsistencies between the model and these data would point to the limitations of single-circuit attractor models of WM storage, to the computational requirements of distributed storage, and by extension, to the role of PPC in distributed storage. Next, we did the same thing with a hierarchical model of PPC and PFC, reasoning that the values of biophysical parameters required to account for the data would identify specific mechanisms and principles for the implementation of distributed storage. If so, then violation of these principles should account for behavioural data from low-performing subjects. Thus, we evaluated our hierarchical model for its ability to account for capacity and overload among subject groups distinguished according to these measures. Finally, we sought to make predictions to test our hypothesis experimentally, approximating electroencephalogram (EEG) recordings over PPC and PFC. We reasoned that if this approximation could account for the different EEG profiles of high- and low-performing subject groups during the storage of memoranda (Fukuda, Woodman, & Vogel, 2015), then its profile during stimulus encoding would be a testable prediction for our hypothesis.

## 2 Methods

Our local-circuit PPC model (the PPC-only model, Figure 1A) is a network of simulated pyramidal neurons and inhibitory interneurons, connected by AMPA, NMDA and GABA receptor conductance synapses (AMPAR, NM-DAR and GABAR). Our hierarchical model (the PPC-PFC model, Figure 1D) is comprised of two such networks, bidirectionally connected. In both models, intrinsic synaptic connectivity within and between classes of neuron was structured according to *in vitro* data, as was the connectivity between networks in the PPC-PFC model. Our chosen parameters and their values are justified in Section 2.6. Note that the synaptic resolution of the models allowed us to approximate EEG signals over PPC and PFC (Results section 3.2.5).

**Figure 1:**
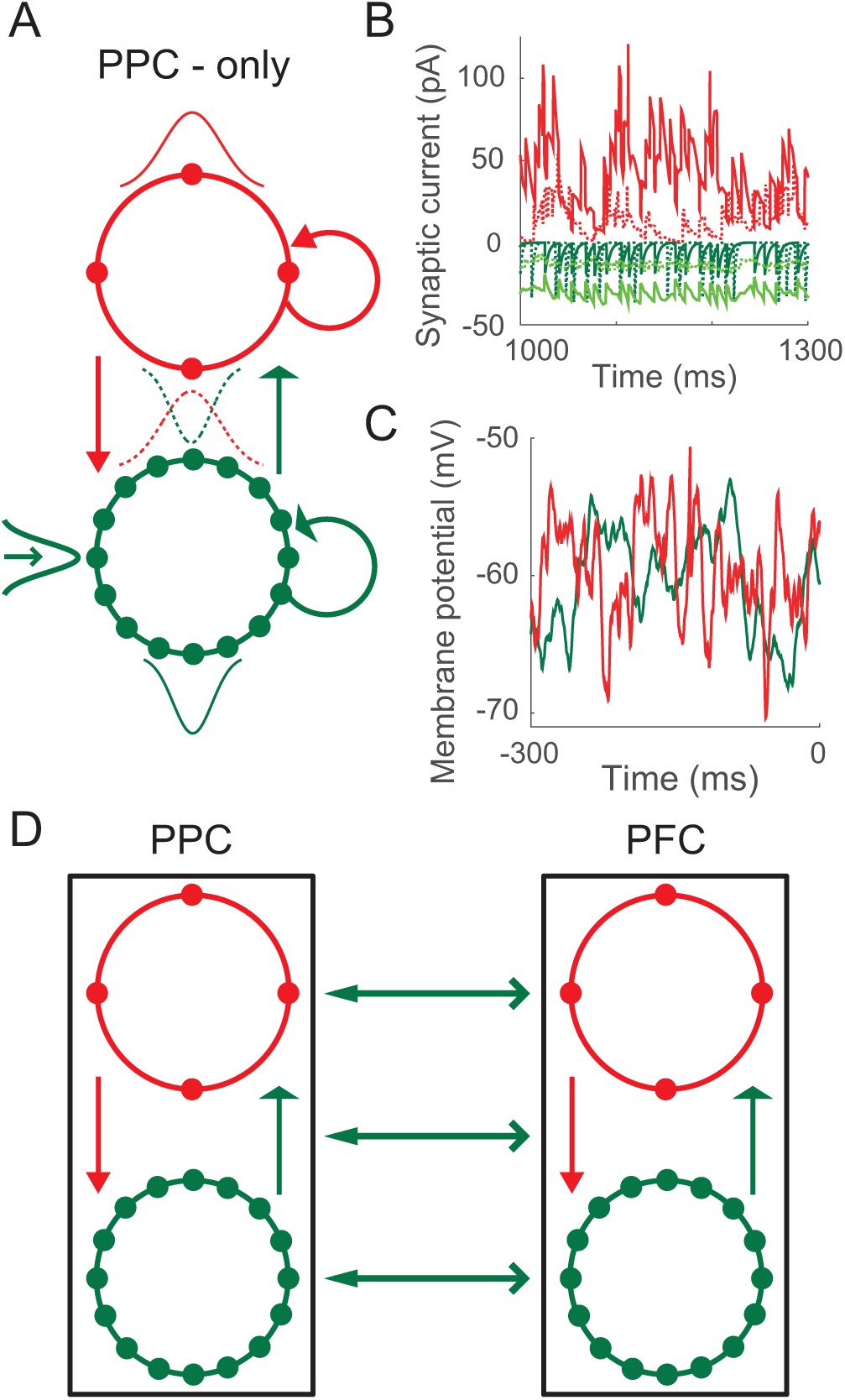
*(preceding page)*: (A) Schematic of the PPC-only model. Solid circles depict pyramidal neurons (green) and inhibitory interneurons (red), arranged periodically by their connectivity structures. The 4-to-1 ratio of pyramidal neurons to interneurons preserves their population sizes in the model. Arced and straight arrows depict synaptic connectivity within and between classes of neuron respectively. Thin Gaussian curves depict the structure of this connectivity (within, solid; between, dotted). The Gaussian curve on the left depicts the RF of a pyramidal neuron. Red, open green and wide green arrows depict GABAR, AMPAR-only, and AMPAR-NMDAR synapses respectively. (B) Synaptic currents onto a pyramidal neuron (solid) and an interneuron (dotted) during the delay interval of the 1-item memory task. Red, light green and dark green curves show GABAR, AMPAR and NMDAR currents respectively. (C) Membrane potential of a pyramidal neuron and an interneuron during the pre-trial interval. (D) Schematic of the PPC-PFC model. The PPC network is identical to the PPC-only model. The PFC network differs only in the strength of GABAR conductance onto pyramidal neurons. Open and thin arrows depict topographically aligned feed-forward and feedback projections, mediated by AMPARs and NMDARs respectively. See Section 2 for description of the model.

We ran simulations of a visuospatial WM task with both models, where the number of items for retention ranged from 1 to 8. On each trial of the task, a stimulus interval was preceded by a pre-trial interval and followed by a delay of 1s. In both models, the items were provided to the PPC network during the stimulus interval and their accurate retention (or otherwise) was determined from its activity at the end of the delay (Sections 2.4 and 2.5).

### 2.1 The network model

Each local circuit is a fully connected network of leaky integrate-and-fire neurons (Tuckwell, 1988), comprised of *N*^*p*^ = 400 simulated pyramidal neurons and *N*^*i*^ = 100 fast-spiking inhibitory interneurons. Each model neuron is described by

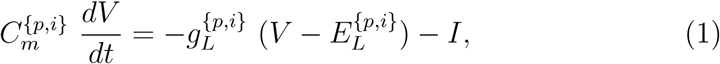

where *C*_*m*_ is the membrane capacitance of the neuron, *g*_*L*_ is the leakage conductance, *V* is the membrane potential, *E*_*L*_ is the equilibrium potential, and *I* is the total input current. When *V* reaches a threshold *ϑ*_*v*_, it is reset to *V*_*res*_, after which it is unresponsive to its input for an absolute refractory period of *τ*_*ref*_. Here and below, superscripts *p* and *i* refer to pyramidal neurons and interneurons respectively, indicating that parameter values are assigned separately to each class of neuron.

The total input current at each neuron is given by

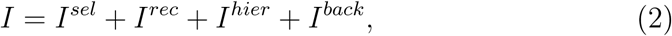

where *I*^*sel*^ is stimulus-selective synaptic current (set to 0 for all neurons in the PFC network and for interneurons in the PPC network), *I*^*rec*^ is recurrent (intrinsic) synaptic current, *I*^*hier*^ is hierarchical (inter-areal) synaptic current projected to PPC from PFC and *vice versa* (set to 0 in single-circuit simulations) and *I*^*back*^ is background current. Of these currents, *I*^*sel*^, *I*^*rec*^ and *I*^*hier*^ are comprised of synaptic currents, and *I*^*back*^ is comprised of synaptic current and injected current. Synaptic currents driven by pyramidal neuron spiking are mediated by simulated AMPA receptor (AMPAR) and/or NMDA receptor (NMDAR) conductances, and synaptic currents driven by interneuron spiking are mediated by simulated GABA receptor (GABAR) conductances. For AMPAR and GABAR currents, synaptic activation (the proportion of open channels) is defined by

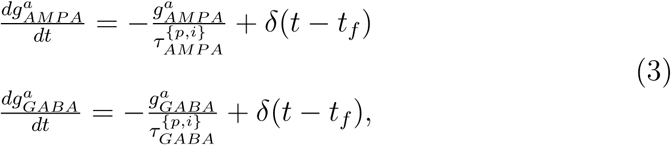

where *τ*_*AMP* *A*_ and *τ*_*GABA*_ are the time constants of AMPAR and GABAR deactivation respectively, *δ* is the Dirac delta function, *t*_*f*_ is the time of firing of a pre-synaptic neuron and superscript *a* indicates that synapses are activated by different sources of spiking activity (selective, recurrent, hierarchical and background). NMDAR activation has a slower rise and decay and is described by

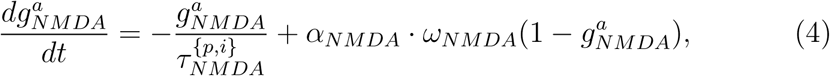

where *τ*_*NMDA*_ is the time constant of receptor deactivation and *α*_*NMDA*_ controls the saturation of NMDAR channels at high pre-synaptic spike frequencies. The slower opening of NMDAR channels is captured by

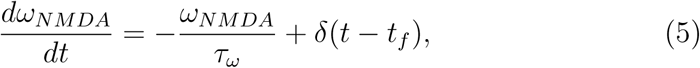

where *τ*_*ω*_ and *η* determine the rate of channel opening and the voltage-dependence of NMDARs respectively.

Recurrent synaptic current to each neuron *j* is defined by

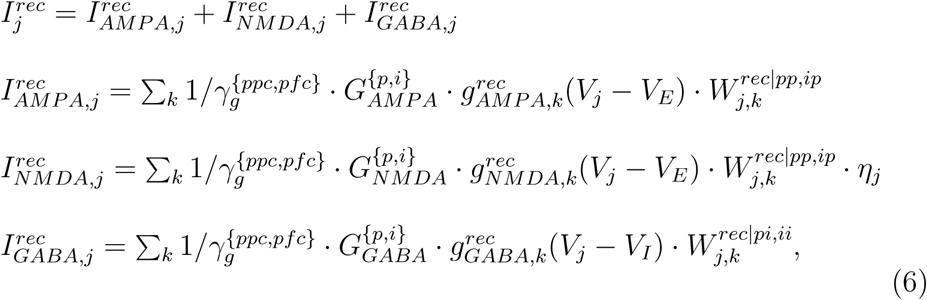

where 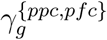 is a scale factor controlling the relative strength of extrinsic and intrinsic synaptic conductance (subscripts *ppc* and *pfc* indicate that its value is assigned separately to each network, see Section 2.6); *G*_*AMPA*_, *G*_*NMDA*_ and *G*_*GABA*_ are the respective strengths of AMPAR, NMDAR and GABAR conductance; *V*_*E*_ is the reversal potential for AMPARs and NMDARs, and *V*_*I*_ is the reversal potential for GABARs; 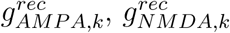 and 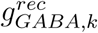 are the activation of AMPAR, NMDAR and GABAR receptors respectively by pre-synaptic neurons *k*; and matrices *W* ^*rec*|*pp,ip*^ and *W* ^*rec*|*pi,ii*^ scale conductance strength or *weight* according to the connectivity structure of the network. This structure depends on the class of neuron receiving and projecting spiking activity, where superscripts *pp, ip, pi* and *ii* denote connections to pyramidal neurons from pyramidal neurons, to interneurons from pyramidal neurons, to pyramidal neurons from interneurons, and to interneurons from interneurons respectively. For each of these structures *s* ∈ {*pp, ip, pi, ii*}, *W* ^*rec*|*s*^ is a Gaussian function of the distance between periodically-arranged neurons, where the weight 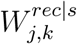 to neuron *j* from neuron *k* is given by

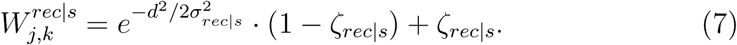

The distance between neurons is defined by *d* = *min*(|*j*-*k*|Δ*x,* 2*π*-|*j*-*k*|Δ*x*) for *W* ^*rec*|*pp*^ and *W* ^*rec*|*ii*^, and by *d* = *min*(|*j* - *z*|Δ*x,* 2*π* -|*j* - *z*|Δ*x*) for *W* ^*rec*|*ip*^ and *W* ^*rec*|*pi*^, where *z* = *N*^*p*^*/N*^*i*^ · *k* for *W* ^*rec*|*ip*^ and *z* = *N*^*i*^*/N*^*p*^ · *k* for *W* ^*rec*|*pi*^. Δ*x* = 2*π/N*^{*p,i*}^ is a scale factor and *σ*_*rec*|*s*_ determines the spatial extent of connectivity. Parameter *ζ*_*rec*|*s*_ allows the inclusion of a baseline weight, with the function normalized to a maximum of 1 (0 ≤ *ζ*_*rec*|*s*_ < 1).

### 2.2 Background activity

For each neuron, *in vivo* cortical background activity is simulated by current *I*^*back*^, defined by

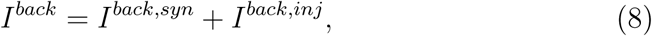

where *I*^*back,syn*^ is driven by synaptic bombardment and *I*^*back,inj*^ is noisy current injection. The former is generated by AMPAR synaptic activation, where independent, homogeneous Poisson spike trains are provided to all neurons at rate *µ*_*back*_. *I*^*back,syn*^ is therefore defined by

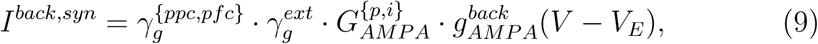

where 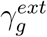 is a scale factor and 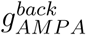 is given in Equation 3.

For *I*^*back,inj*^, we used the point-conductance model by (Destexhe, Rudolph, Fellous, & Sejnowski, 2001):

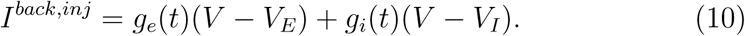

The time-dependent excitatory and inhibitory conductances *g*_*e*_(*t*) and *g*_*i*_(*t*) are updated at each timestep Δ*t* according to

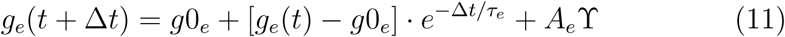

and

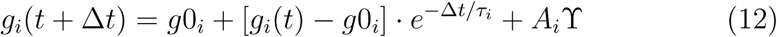

respectively, where *g*0_*e*_ and *g*0_*i*_ are average conductances, *τ*_*e*_ and *τ*_*i*_ are time constants, and ϒ is normally distributed random noise with 0 mean and unit standard deviation. Amplitude coefficients *A*_*e*_ and *A*_*i*_ are defined by

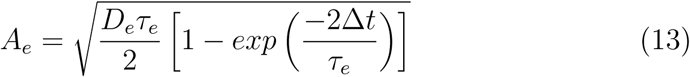

and

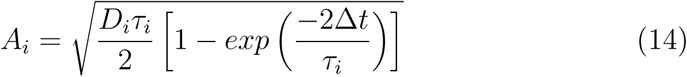

respectively, where 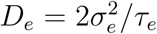 and 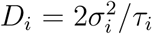 are noise ‘diffusion’ coefficients. See (Destexhe et al., 2001) for the derivation of these equations.

### 2.3 The PPC-PFC model

In the hierarchical model, inter-areal projections mediate synaptic currents *I*^*hier*^ ∈ {*I*^*ff*^, *I*^*fb*^}, where superscript *ff* (*fb*) refers to feedforward (feedback) currents onto neurons in the PFC (PPC) network from neurons in the PPC (PFC) network. Only pyramidal neurons make inter-areal projections, where feedforward projections are mediated by AMPARs (onto pyramidal neurons only) and feedback projections are mediated by NMDARs (onto pyramidal neurons and interneurons). Feed-forward currents at each pyramidal neuron *j* in the PFC network are defined by

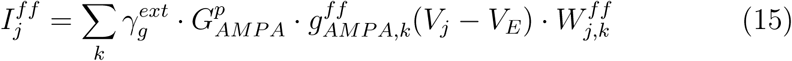

where 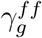 is a scale factor, 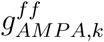 is the activation of AMPAR receptors by pre-synaptic pyramidal neurons *k* in the PPC network, and matrix 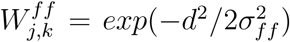 scales conductance strength according to the structure of FF connectivity. Constant *d* is given above for recurrent synaptic structure *W* ^*rec*|*pp*^, where the two networks are topographically aligned, *i.e.* the lateral distance between neuron *j* in the PFC network and neuron *k* in the PPC network is the same as that between neurons *j* and *k* within either network.

Feedback currents at each neuron *j* in the PPC network are defined by

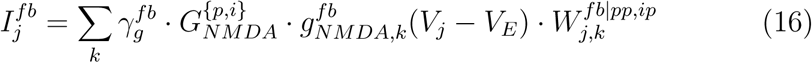

where 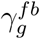 is a scale factor, 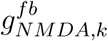 is the activation of NMDAR receptors by pre-synaptic pyramidal neurons *k* in PFC, and matrices *W* ^*fb*|*pp,ip*^ scale conductance strength according to the structure of FB connectivity. Each of these structures *s* ∈ {*pp, ip*} is defined by 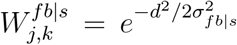, where *d* is defined for *W* ^*fb*|*pp*^ and *W* ^*fb*|*ip*^ in the same way as for *W* ^*rec*|*pp*^ and *W* ^*rec*|*ip*^ respectively above.

### 2.4 Simulated working memory task

We simulated the stimulus array by providing independent, homogeneous Poisson spike trains to all pyramidal neurons *j* in the PPC network, where spike rates were drawn from a normal distribution with mean *µ*_*sel*_ corresponding to the centre of a Gaussian response field (RF) defined by 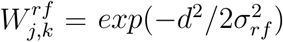. Constant *d* is given above for recurrent synaptic structure *W* ^*rec*|*pp*^, *σ*_*rf*_ determines the width of the RF and subscript *k* indexes the neuron at the RF centre. Spike response adaptation by upstream visually responsive neurons was modelled by a step-and-decay function

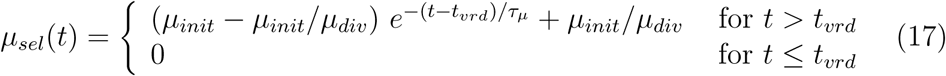

where *µ*_*init*_ determines the initial spike rate, *µ*_*div*_ determines the asymptotic rate, *τ*_*µ*_ determines the rate of upstream response adaptation, and *t*_*vrd*_ is a visual response delay. These selective spike trains were provided for 300ms, following the 300ms pre-trial interval and followed by a 1000ms delay (*e.g.* Figure 2A). The stimuli were mediated by AMPARs only, so for all pyramidal neurons *j* in the PPC network,

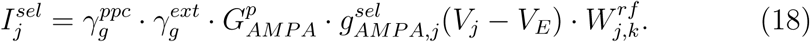

**Figure 2:**
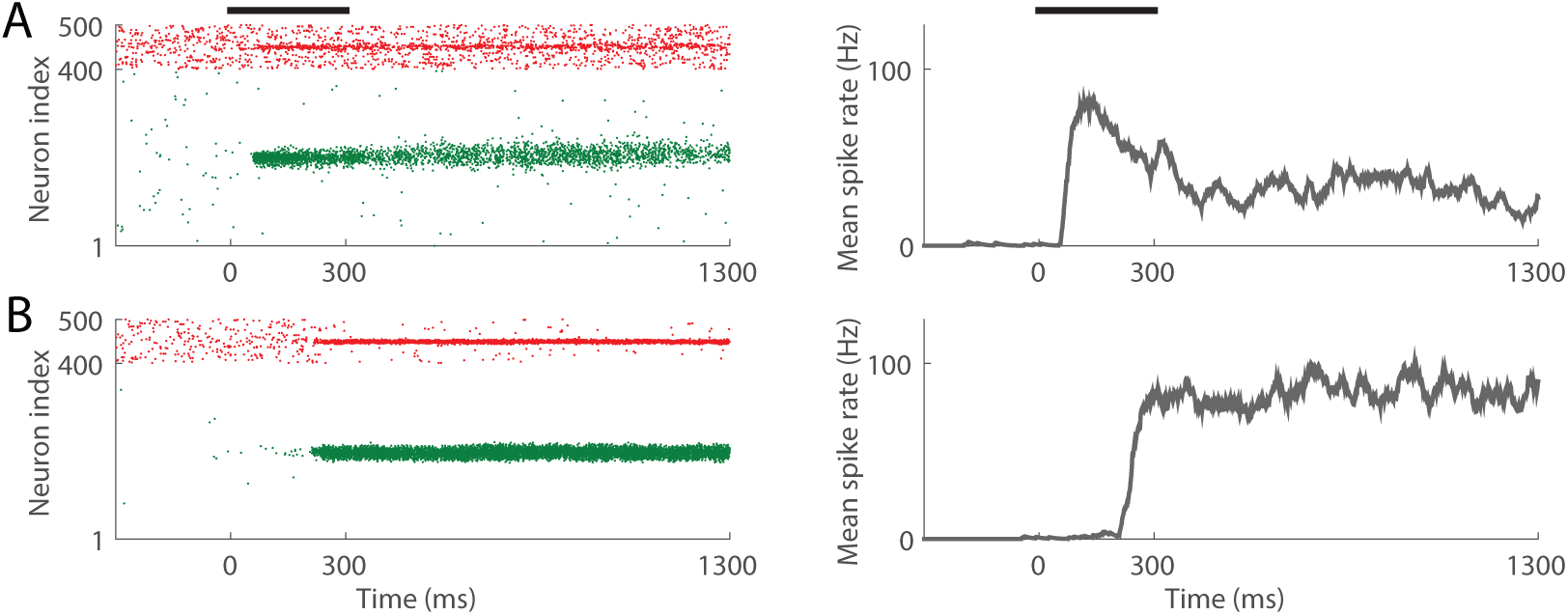
A single trial of the 1-item memory task with the PPC-only model for the lowest and highest values of control parameter 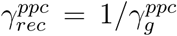, furnishing weak (A, 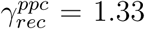) and strong (B, 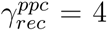) recurrent dynamics respectively. In raster plots, pyramidal neurons and interneurons are indexed from 1 - 400 and 401 - 500 respectively. Each right-side panel shows the mean rate of the item-encoding pyramidal population. Thick horizontal bars show the timing of the stimulus.

All simulations were run with the standard implementation of Euler’s forward method, where the timestep was Δ*t* = 0.25ms.

### 2.5 Determining working memory performance

We ran 100 trials with 1 - 8 stimuli (henceforth the *n*-item memory task; 1 ≤ *n* ≤ 8). To determine WM performance on each trial, spike density functions (SDFs) were calculated for all pyramidal neurons in the network by convolving their spike trains with a rise-and-decay function

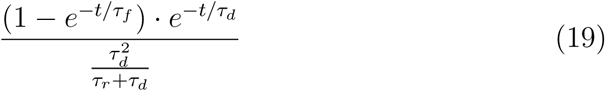

where *t* is the time following stimulus onset and *τ*_*r*_ = 1ms and *τ*_*d*_ = 20ms are the time constants of rise and decay respectively (Thompson, Hanes, Bichot, & Schall, 1996; Standage & Paré, 2011). On each *n*-item trial, we calculated the mean of the SDFs over the last 300ms of the delay, obtaining the average activity over the network, and then partitioned the network into *n* equal regions. The location of each item was centred within each region. We then fit the mean activity in each region with a Gaussian function with four parameters: the height of the peak, the position of the peak, the standard deviation (controlling width), and the height that is approached asymptotically from the peak. An item was considered accurately stored if the fitted Gaussian satisfied three criteria: the height parameter *h* exceeded 30Hz, the difference between *h* and the fitted asymptote on both sides of the peak exceeded *h/*2Hz, and the position parameter was within Δ*c* = 10 degrees of the centre of the RF for that item. For the first criterion, we chose 30Hz because in electrophysiological experiments with macaque monkeys (Johnston et al, SfN abstracts, 2009), memory trials were discarded if the recorded PPC neuron did not fire at least 10 spikes during the last 300ms of the delay (10*/*0.3s ≈ 30Hz). The second criterion dictates that items are only considered accurately stored if the population response is discriminable. The third criterion ensures that the memory of the location of the item is close to the actual location.

### 2.6 Parameter values

In setting parameter values in the two models, our aim was to justify every value by anatomical and physiological data, thus constraining our choices as much as possible, and then to use control parameters to explore the models’ performance on simulated WM tasks. In the PPC-only model, our control parameter was 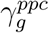 (Equations 6, 9 and 18), governing the relative strengths of extrinsic and intrinsic synaptic conductance and therefore the strength of recurrent processing (see Section 3.1). Parameter values for the PPC-only network are provided in Table 1 and justified by Standage and Paré (2018).

**Table 1:**
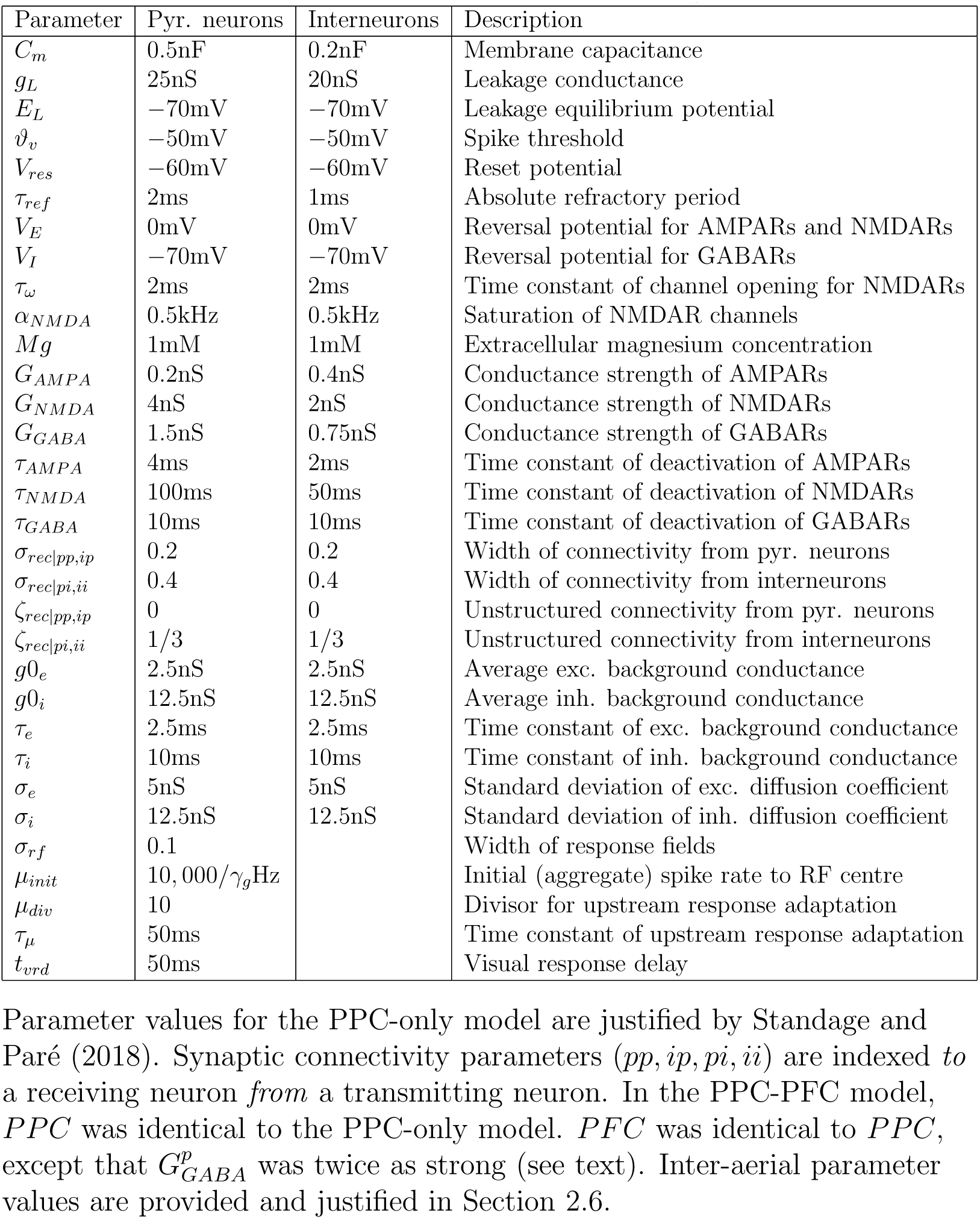
Parameters of the PPC-only model

| Parameter | Pyr. neurons | Interneurons | Description |
| --- | --- | --- | --- |
| $C_m$ | 0.5nF | 0.2nF | Membrane capacitance |
| $g_L$ | 25nS | 20nS | Leakage conductance |
| $E_L$ | -70mV | -70mV | Leakage equilibrium potential |
| $\vartheta_v$ | -50mV | -50mV | Spike threshold |
| $V_{res}$ | -60mV | -60mV | Reset potential |
| $\tau_{ref}$ | 2ms | 1ms | Absolute refractory period |
| $V_E$ | 0mV | 0mV | Reversal potential for AMPARs and NMDARs |
| $V_I$ | -70mV | -70mV | Reversal potential for GABARs |
| $\tau_\omega$ | 2ms | 2ms | Time constant of channel opening for NMDARs |
| $\alpha_{NMDA}$ | 0.5kHz | 0.5kHz | Saturation of NMDAR channels |
| $Mg$ | 1mM | 1mM | Extracellular magnesium concentration |
| $G_{AMPA}$ | 0.2nS | 0.4nS | Conductance strength of AMPARs |
| $G_{NMDA}$ | 4nS | 2nS | Conductance strength of NMDARs |
| $G_{GABA}$ | 1.5nS | 0.75nS | Conductance strength of GABARs |
| $\tau_{AMPA}$ | 4ms | 2ms | Time constant of deactivation of AMPARs |
| $\tau_{NMDA}$ | 100ms | 50ms | Time constant of deactivation of NMDARs |
| $\tau_{GABA}$ | 10ms | 10ms | Time constant of deactivation of GABARs |
| $\sigma_{rec\vert pp,ip}$ | 0.2 | 0.2 | Width of connectivity from pyr. neurons |
| $\sigma_{rec\vert pi,ii}$ | 0.4 | 0.4 | Width of connectivity from interneurons |
| $\zeta_{rec\vert pp,ip}$ | 0 | 0 | Unstructured connectivity from pyr. neurons |
| $\zeta_{rec\vert pi,ii}$ | 1/3 | 1/3 | Unstructured connectivity from interneurons |
| $g_{0e}$ | 2.5nS | 2.5nS | Average exc. background conductance |
| $g_{0i}$ | 12.5nS | 12.5nS | Average inh. background conductance |
| $\tau_e$ | 2.5ms | 2.5ms | Time constant of exc. background conductance |
| $\tau_i$ | 10ms | 10ms | Time constant of inh. background conductance |
| $\sigma_e$ | 5nS | 5nS | Standard deviation of exc. diffusion coefficient |
| $\sigma_i$ | 12.5nS | 12.5nS | Standard deviation of inh. diffusion coefficient |
| $\sigma_{rf}$ | 0.1 | | Width of response fields |
| $\mu_{init}$ | 10,000/$\gamma_g$Hz | | Initial (aggregate) spike rate to RF centre |
| $\mu_{div}$ | 10 | | Divisor for upstream response adaptation |
| $\tau_\mu$ | 50ms | | Time constant of upstream response adaptation |
| $t_{vrd}$ | 50ms | | Visual response delay |

Parameter values for the PPC-only model are justified by Standage and Paré (2018). Synaptic connectivity parameters ( $pp, ip, pi, ii$ ) are indexed *to* a receiving neuron *from* a transmitting neuron. In the PPC-PFC model, *PPC* was identical to the PPC-only model. *PFC* was identical to *PPC*, except that $G_{GABA}^p$ was twice as strong (see text). Inter-aerial parameter values are provided and justified in Section 2.6.

In the PPC-PFC model, the PPC network was identical to the PPC-only model. The PFC network was identical to the PPC network, except that GABAR conductance onto pyramidal neurons was twice as strong 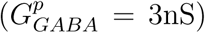, facilitating strong competition during stimulus encoding. We limited the number of parameters in the PPC-PFC model by making three simplifying assumptions. Firstly, we assumed that only pyramidal neurons make inter-areal projections [see Jones (1984); White (1989); Tomioka and Rockland (2007)] and we therefore omitted inter-areal projections from inhibitory interneurons. Secondly, based on evidence that feedforward and feedback excitation are predominantly mediated by AMPARs and NMDARs respectively (Self, Kooijmans, Supéer, Lamme, & Roelfsema, 2012), we omitted feedforward NMDARs and feedback AMPARs altogether. Thirdly, feed-forward and feedback projections were each assumed to be topographically organized, based on evidence that the proportion of feedforward supragranular projections and feedback infragranular projections increase with hierarchical distance between cortical areas. Since supragranular and infragranular projections are believed to be topographic and diffuse respectively, and since PPC and lateral prefrontal cortex are separated by a single hierarchical layer [see Goldman-Rakic (1988)], we assumed that the proportion of topographic supragranular projections and diffuse infragranular projections between these areas is approximately equal in both directions. See Markov and Kennedy (2013) for a more thorough description of these macrocircuit principles. Given the diffuse activity already simulated by background current *I*^*back*^ (Equations 9 and 10), we ignored the effect of infragranular feed-forward and feedback projections between the PPC and PFC networks, and simulated topographic supragranular connectivity only. The width of feed-forward RFs (PFC network) was the same as for stimulus-selective RFs in the PPC network (*σ*_*ff*_ = *σ*_*rf*_; Equations 15 and 18), based on evidence that differences in dendritic branching are minimal between hierarchically adjacent cortical areas (Elston, 2002). Feedback RFs (PPC) were wider than feedforward RFs (*σ*_*fb*_ = 1.5 · *σ*_*ff*_, Equations 16 and 15) because dendritic branching in supragranular layers is more extensive than in (feedforward input) layer 4. For simplicity, we omitted FF projections onto PFC interneurons, having omitted stimulus-selective inputs to PPC interneurons [a common approach with local-circuit models of this class, *e.g.* Compte, Brunel, Goldman-Rakic, and Wang (2000); Wei et al. (2012); Standage and Paré (2018)]. Control parameters for the PPC-PFC model are described in Section 3.2.

## 3 Results

### 3.1 Competitive encoding in the PPC-only model alleviates overload, but conflicts with experimental data

In the PPC-only model, we used control parameter 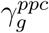 (Equations 6, 9 and 18 in Methods) to modulate network dynamics, scaling the strength of all extrinsic synapses inversely with that of all intrinsic synapses. By applying this scale factor to excitatory and inhibitory synapses onto pyramidal neurons and interneurons, we were able to maintain excitatory/inhibitory balance, controlling retention ability by recurrent excitation and competitive interactions by lateral inhibition. We determined the floor and ceiling on 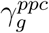 by stipulating that the model must perform the 1-item memory task with 90% accuracy. To this end, we ran 100 trials of the task for a range of values of 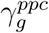 (increments of 0.05), finding that our criterion was satisfied for 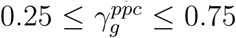. Figure 2 shows example trials of the 1-item task for the highest and lowest values of this parameter. Because lower values support stronger recurrent dynamics (intrinsic synaptic conductance values are divided by 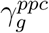, Equation 6), it is intuitive to define parameter 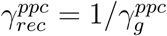, so that higher values support stronger recurrent dynamics. We use the latter term in the description of our results below.

We ran 100 trials for each value of 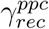 in each load condition of the multiple-item memory task (2 ≤ *n* ≤ 8, where *n* is the number of items). See Figure 4 for example trials of the 8-item task. For each load condition, we calculated the mean number of items accurately encoded during the stimulus interval, referred to as the effective load *E*(*n*), and the mean number of items accurately retained over the memory delay, referred to as capacity *K*(*n*). We refer to the maximum value of *K*(*n*) as peak capacity 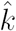 and we define WM overload as Θ = 1-*K*[max[(*n*)]/max[*K*(*n*)], *i.e.* overload refers to a decrease in capacity on the 8-item task, relative to peak capacity. For convenience, these and other terms are defined in Table 2.

**Table 2:** Summary of terminology

| Term | Description |
| --- | --- |
| PPC | Posterior parietal cortex |
| PFC | Lateral prefrontal cortex |
| PPC | The PPC network in the PPC-only model and the PPC-PFC model |
| PFC | The PFC network in the PPC-PFC model |
| $\gamma_g^{ppc}$ | Control param. for *PPC*, scaling syn. conductance (Eq. 6, 9 and 18) |
| $\gamma_g^{pfc}$ | Control param. for *PFC*, scaling syn. conductance (Eq. 6, 9 and 18) |
| $\gamma_g^{fb}$ | Control param. scaling conductance at feedback synapses to *PPC* from *PFC* (Eq. 16) |
| PPC modulation | Modulation of recurrent dynamics in *PPC* by $\gamma_{rec}^{ppc} = 1/\gamma_g^{ppc}$ |
| PFC modulation | Modulation of recurrent dynamics in *PPC* by $\gamma_{rec}^{ppc} = 1/\gamma_g^{ppc}$ |
| Inter-aerial modulation | Modulation of feedback connectivity by $\gamma_g^{fb}$ |
| 2-parameter modulation | Simultaneously varying two control parameters, while holding the third one fixed |
| 3-parameter modulation | Simultaneously varying all three control parameters |
| Memory load $n$ | Number of WM items on a given trial, a.k.a the load condition |
| Capacity $K(n)$ | Mean number of items accurately retained over the memory delay for load $n$ |
| Peak capacity $\hat{k}$ | Maximum value of $K(n)$ over all $n$ |
| Effective load $E(n)$ | Mean number of items accurately encoded during the stimulus interval for load $n$ |
| $E'$ | Minimum ratio of effective load to actual load over all $n$, *i.e.* $E' = \min[E(n)/n]$ |
| WM overload $\Theta$ | $\Theta = 1 - K[\max(n)]/\max[K(n)]$ |

Summary of terminology defined in the text, including control parameters for the PPC-only model and the PPC-PFC model.

**Figure 3:**
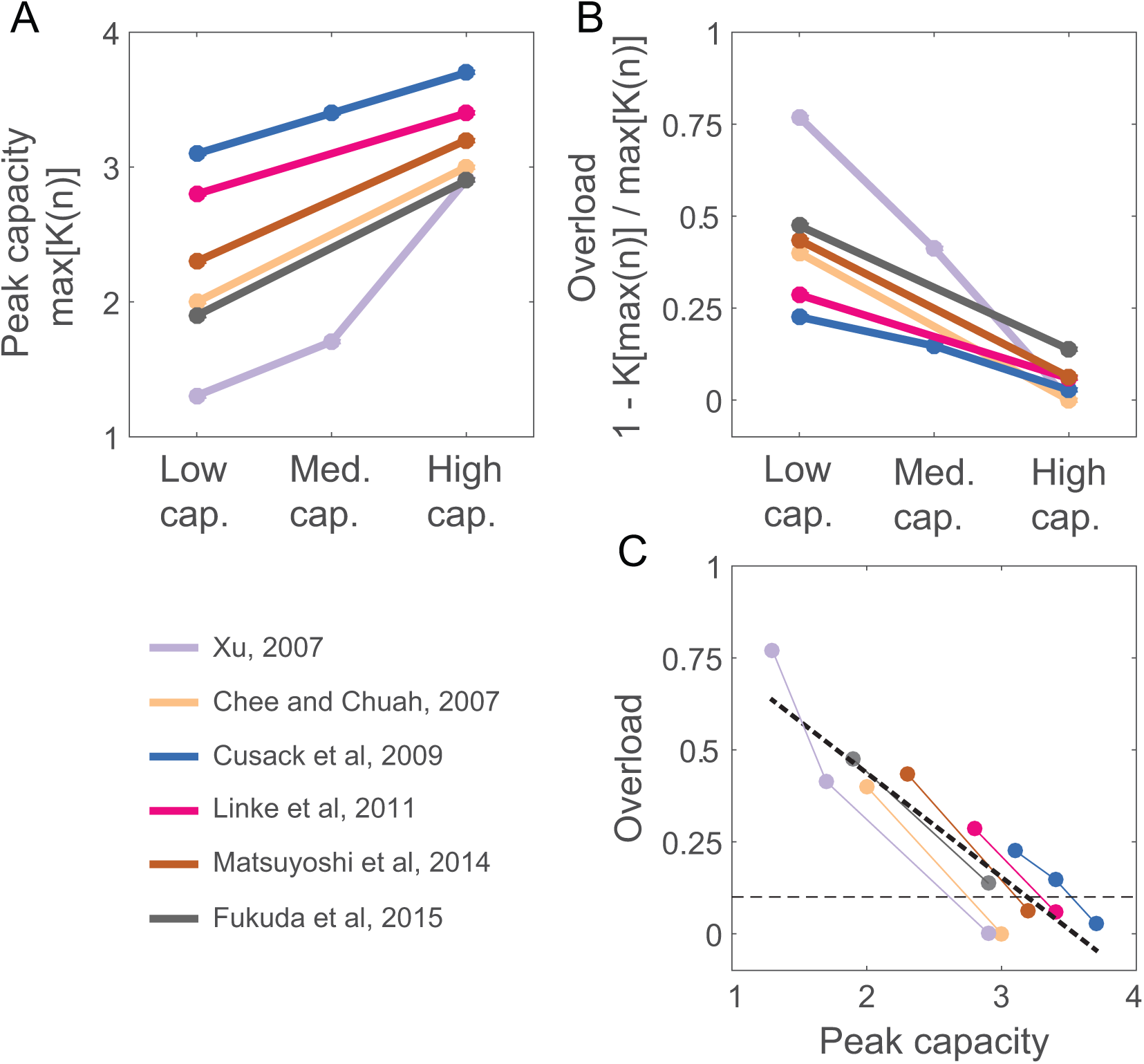
A summary of behavioural data showing WM overload. (A-B) Peak capacity (A) and overload (B) shown by six studies of multiple-item WM. More pronounced overload occurs with lower-capacity performance (across subjects, tasks and conditions of the same task; see text), such that peak capacity and overload show a strong negative correlation (C). (C) Overload over peak capacity (Pearson correlation coefficient *r* = 0.889, *p* = 2.084*e*-5).

**Figure 4:**
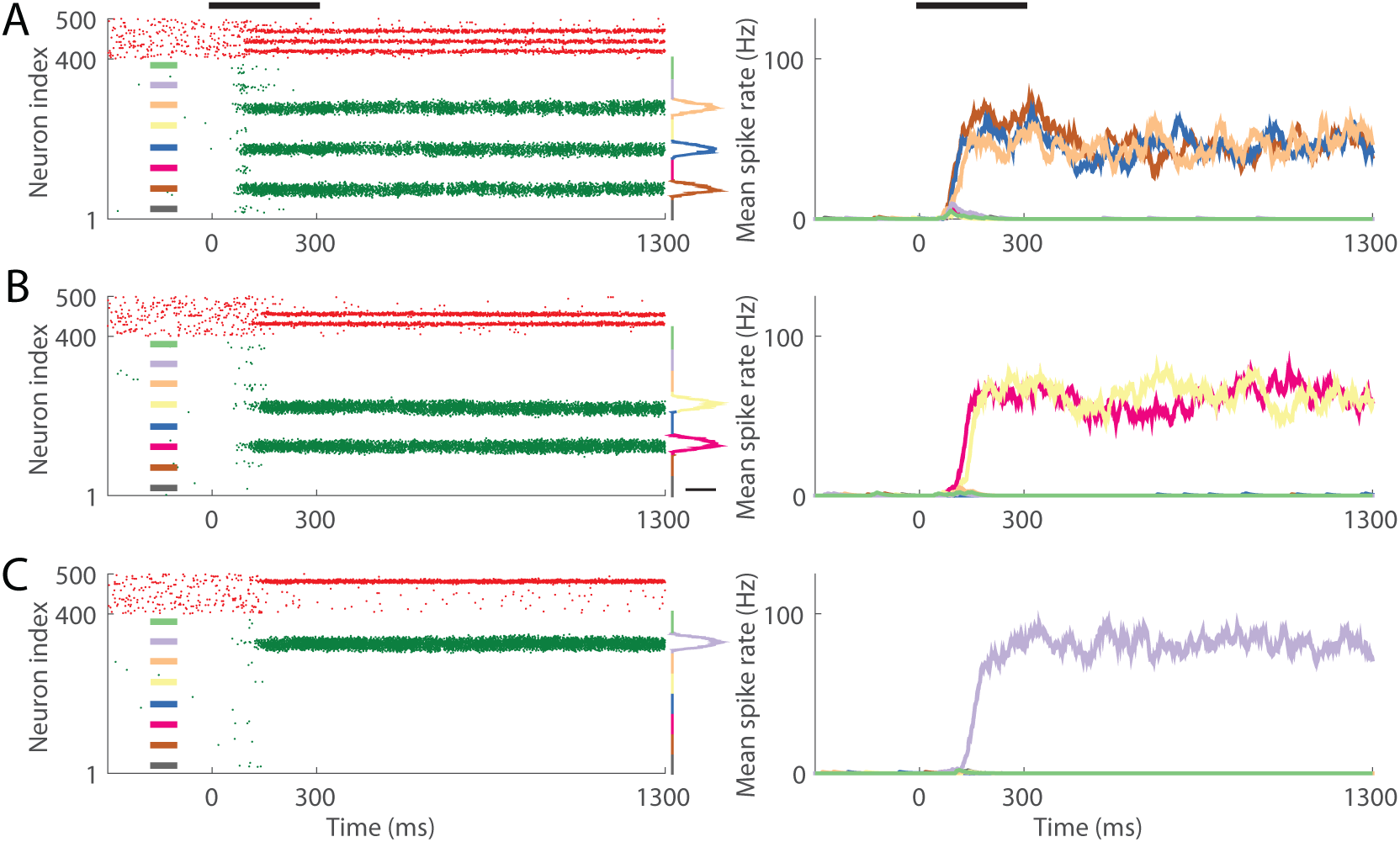
A single trial of the 8-item memory task with the PPC-only model for each of three values of control parameter 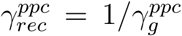, on which the network accurately retained three (A, 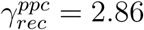), two (B, 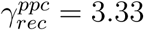) and one (C, 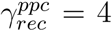) item(s). Raster plots (left) and mean spike rates (right) are described in Figure 2. Here, colours show the correspondence between item-encoding populations in the left and right panels. Competition during the stimulus interval reduces the load to 3 (A), 2 (B) and 1 (C) item(s) by the onset of the memory delay.

Our results with the PPC-only model distinguished between two subsets of our control parameter. For 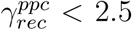, peak capacity increased with the strength of recurrent dynamics (Figure 5A), almost all stimuli were encoded (Figure 5B), and overload was catastrophic for all values of the parameter [*K*(8) was close to 0, Figure 5C]. For 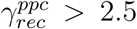, peak capacity, effective load and overload all decreased with stronger recurrent dynamics. Thus, 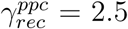 separated two qualitatively different regimes, both of which supported the control of peak capacity by 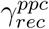, but only one of which supported the control of effective load and overload. Consequently, we limit further consideration of the PPC-only model to 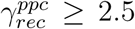, where stronger recurrent dynamics imposed more intense competition during stimulus encoding, limiting effective load and thereby reducing overload. In this regime, peak capacity was 2 or 3 items for all but the strongest recurrent dynamics, consistent with that of human and monkey subjects [*e.g.* Luck and Vogel (1997); Heyselaar, Johnston, and Paré (2011)], but the alleviation of overload came at the expense of peak capacity (significant positive correlation between these measures, Figure 5D). The available data show the opposite trend: overload is typically more pronounced among lower-capacity subjects (Linke et al., 2011; Matsuyoshi et al., 2014; Fukuda, Woodman, & Vogel, 2015), on lower-capacity tasks (Xu, 2007) and in lower-capacity conditions of the same task (Chee & Chuah, 2007). These data are summarised in Figure 3. Furthermore, recurrent dynamics strong enough to eliminate overload altogether furnished a peak capacity of 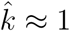 item, unrealistically low for healthy adult subjects ^1^. Thus, the PPC-only model provides proof of concept for competitive encoding, but is qualitatively inconsistent with prominent trends in behavioural data (Figure 3). Furthermore, spiking activity in the model conflicts with neural data from PPC, which typically show a response to all items in a stimulus array, prior to the selection of task-relevant items [*e.g.* Thomas and Paré (2007)]. This discrepancy is implicit in Figure 5B and explicit in Figure 4A-C, where the effective load on the 8-item task is three, two and one items respectively. We therefore turned to the PPC-PFC model, investigating its ability to account for these and other data.

**Figure 5:**
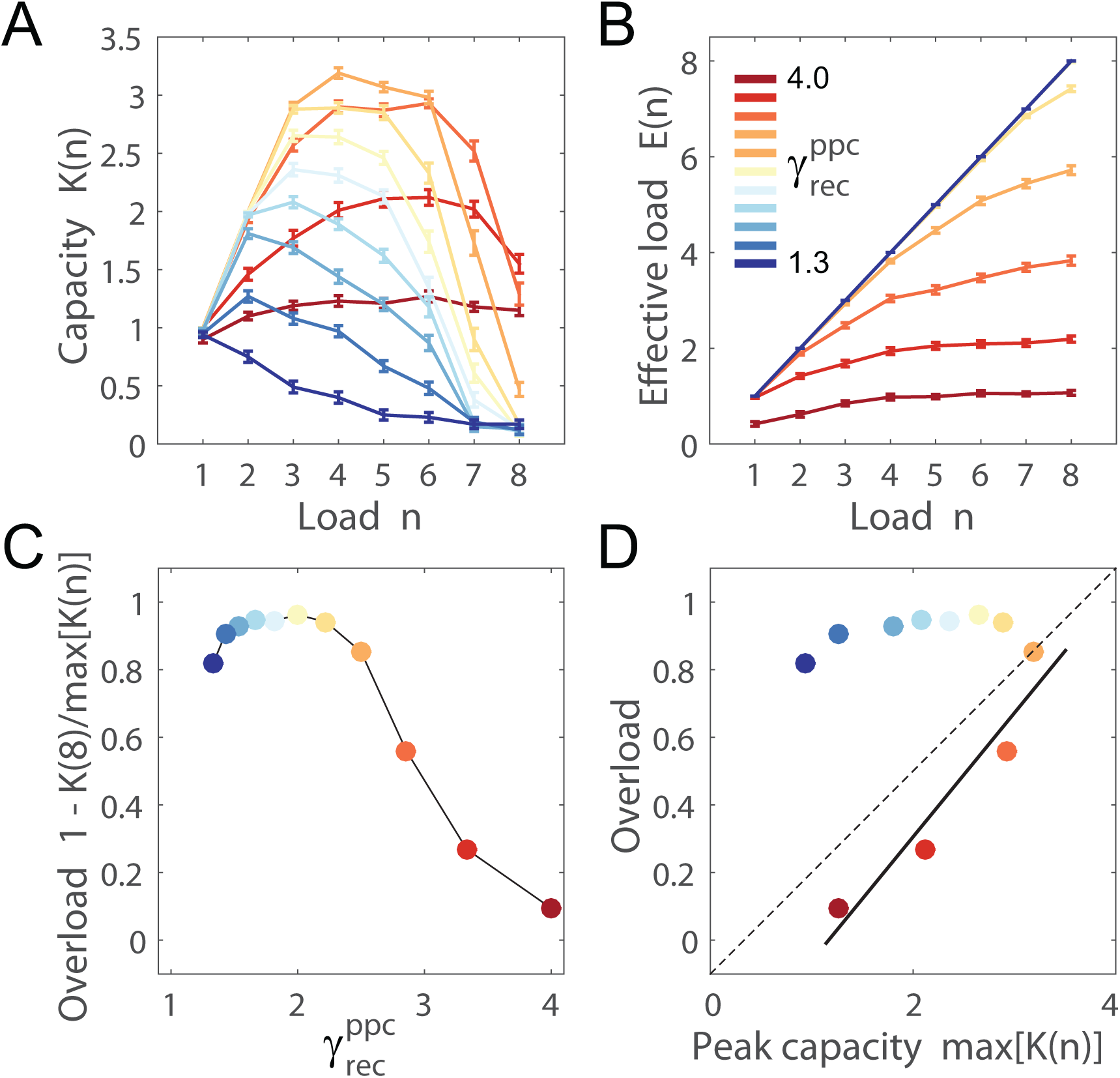
*(preceding page)*: Competitive encoding alleviates overload in the PPC-only model, but at a cost to peak capacity (see text). (A) Capacity *K*(*n*) in each load condition (number of items *n*) for each value of control parameter 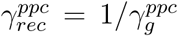. Error bars show standard error. Colour coding interpolates between weakest (dark blue) and strongest (dark red) recurrent dynamics (see legend in panel B). Peak capacity is highest for 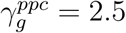 and *K*(8) is close to 0 for 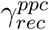 below this value [*K*(8) < 0.15]. Thus, there is no advantage to 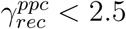. (B) The mean number of items encoded during the stimulus interval for each value of 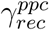, referred to as the effective load *E*(*n*). For 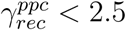, *E*(*n*) > 0.9 · *n* in all load conditions. For 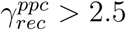, encoding is increasingly competitive (bottom four curves). (C) WM overload for each value of 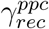, calculated as the relative change from peak capacity to *K*(8) (see text). For 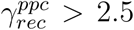, overload is greatly reduced. (D) Overload over peak capacity. For 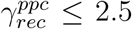, the linear fit (solid line) shows a significant positive correlation between these measures (*r* = 0.955, *p* = 0.045). The dashed line depicts two qualitatively different regimes in the model, separated by 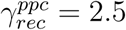.

### 3.2 The PPC-PFC model alleviates overload in a manner consistent with neural and behavioural data

We used three control parameters with the PPC-PFC model. Once again, we controlled recurrent dynamics in the PPC network (henceforth *PPC*) with parameter 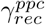. We controlled recurrent dynamics in the PFC network (henceforth *PFC*) in the same way with parameter 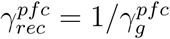 (Equations 6, 9 and 18) and we controlled the strength of feedback projections to *PPC* from *PFC* with parameter 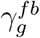 (Equation 16). These control parameters and related terms are defined in Table 2. We began by assigning values to these parameters that explicitly capture the computational principles identified above: weak recurrent dynamics in PPC, allowing the initial encoding of all items, and strong recurrent dynamics in PFC, supporting strong competition during stimulus encoding. Under this approach, persistent mnemonic activity was supported by strong feedback connectivity to *PPC* from *PFC*. An example of the 8-item memory task is shown in Figure 6, where our chosen parameter values are provided in the figure caption. The model had a peak capacity of 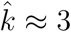 items, overcame overload almost entirely (Θ ≈ 0.075) and allowed the initial encoding of all items in *PPC* (Figure 7). To emphasize the effectiveness of the proposed mechanism, we compared these results to the best possible performance by the PPC-only model, selecting the highest capacity for each load condition across all values of 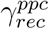. As shown in Figure 7A, the PPC-only model cannot reduce overload without a cost to peak capacity, even if we assume perfect load-dependent modulation of network dynamics by our control parameter.

**Figure 6:**
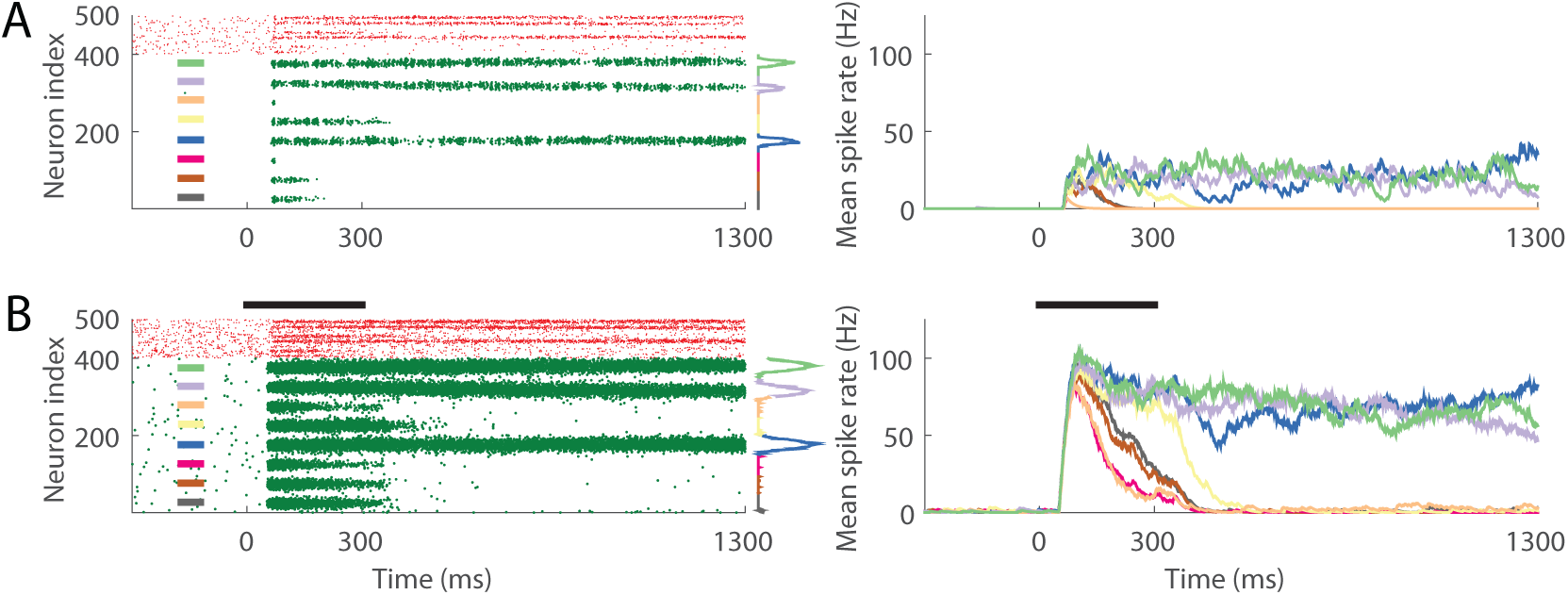
A single trial of the 8-item memory task with the PPC-PFC model, on which three items were accurately retained. Raster plots (right), mean spike rates (left) and corresponding colour schemes are described in Figures 2 and 4. Here, spiking activity is shown in the PPC network (*PPC*, bottom) and the PFC network (*PFC*, top). Weak recurrent dynamics in *PPC* 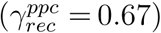 support the initial encoding of all items in the network, but the effective load is reduced by competitive encoding in *PFC* 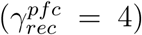. Persistent activity is supported by inter-areal projections 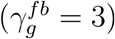.

**Figure 7:**
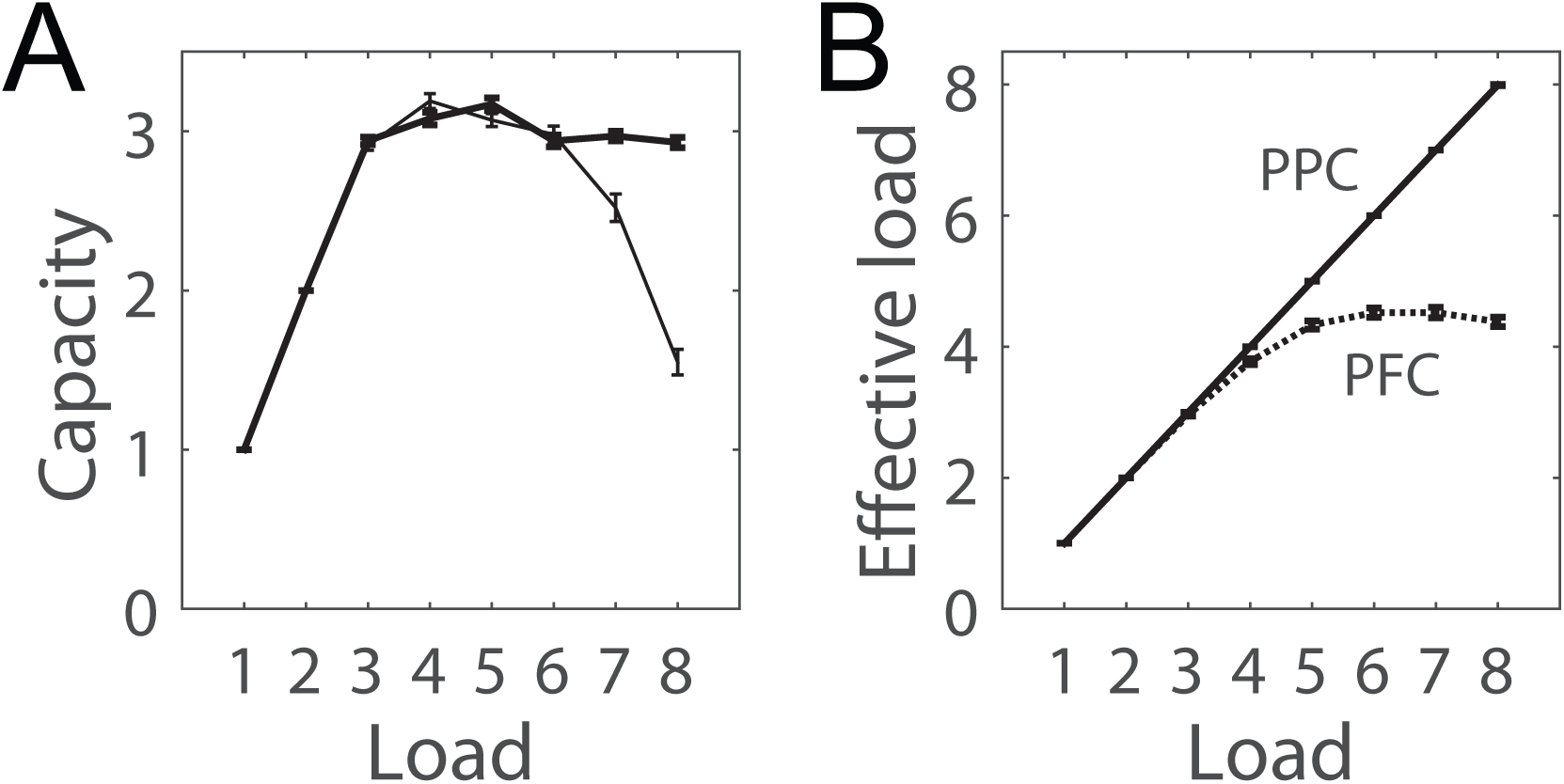
Competitive encoding in *PFC* alleviates overload in the PPC-PFC model, where weak recurrent dynamics in *PPC* permit the initial encoding of all items. (A) Capacity *K*(*n*) for the PPC-PFC model (thicker curve) and the ‘best of’ *K*(*n*) in the PPC-only model [thin curve; highest *K*(*n*) over all 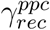, see text]. (B) Effective encoding *E*(*n*) in the PPC-PFC model, where solid and dashed curves correspond to *PPC* and *PFC* respectively. Error bars show standard error. Control parameters are 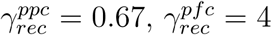 and 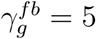.

#### 3.2.1 Simulated cognitive control by modulation of PPC dynamics, PFC dynamics and inter-aerial connectivity

Having determined that hierarchical recruitment of competition alleviates overload in the PPC-PFC model, we sought to determine whether less overload would be seen under parameters supporting higher peak capacity, per the available data (Figure 3) (Chee & Chuah, 2007; Cusack et al., 2009; Matsuyoshi et al., 2014; Fukuda, Woodman, & Vogel, 2015). To this end, we ran a block of trials (100 trials for memory loads 1 : 8) for a range of values of each control parameter, calculating peak capacity 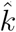 and overload Θ for each combination of values (each configuration). We then determined the correlation (Pearson’s r) between peak capacity and overload 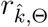 with monotonic changes to our parameter values, simulating cognitive control by modulation of PPC dynamics (varying 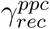, henceforth PPC modulation), PFC dynamics (varying 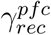, henceforth PFC modulation) and the functional connectivity between PPC and PFC (varying 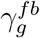, henceforth inter-aerial modulation). Overall, we ran blocks for 500 configurations, spanning 10 values of 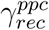 (0.67 : 0.1 : 1.67), five values of 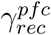 (2.5 : 0.05 : 5) and ten values of 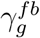 (1 : 1 : 10). This range of values was sufficiently broad and fine-grained for monotonic trends in 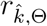 to saturate and for non-monotonic trends to clearly reverse direction (see below).

We first considered our control parameters in isolation from one another, calculating 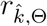 over all values of a given parameter for every combination of the remaining two parameters. PPC modulation yielded a strong negative correlation over a broad region of the 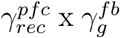 parameter space (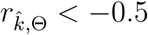 for 14 out of 50 combinations of 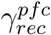 and 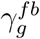), where moderate to strong recurrent dynamics in *PFC* coincided with moderate to strong feedback projections to *PPC* (Figure 8A). Thus, a negative correlation was seen when competitive encoding was strong enough to select a manageable number of items and feedback projections were strong enough to sustain their neural representations over the delay. Notably, peak capacity increased and overload decreased as recurrent dynamics in *PPC* were *weakened* (lower 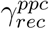, Figure 9A-C), suggesting that if PFC selects memoranda and inter-areal projections sustain their representations after stimulus-offset, then the best thing for PPC to do is give way to its inputs, thereby limiting competition during the delay.

**Figure 8:**
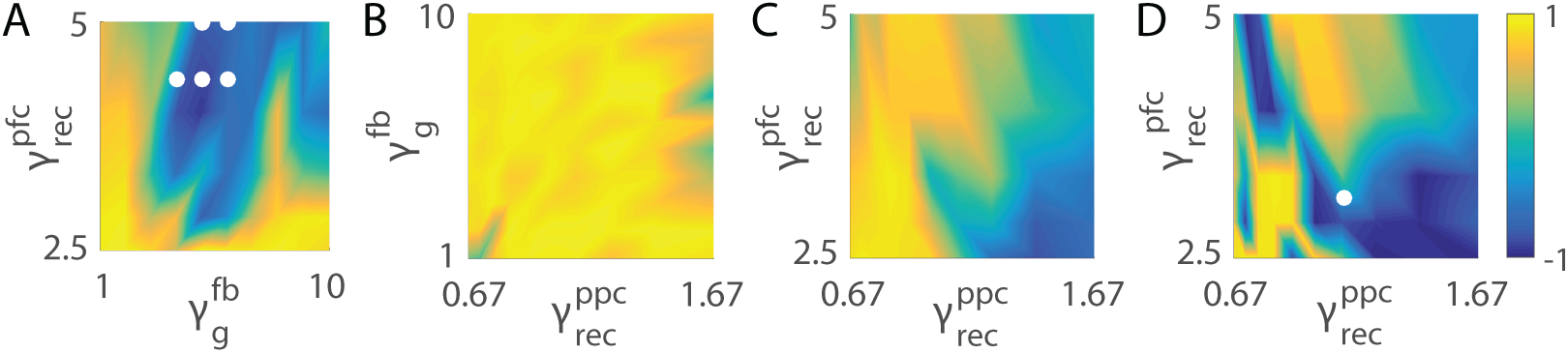
Correlation between peak capacity and overload in the PPC-PFC model under PPC modulation (varying 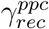, A), PFC modulation (varying 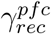, B) and modulation of PPC-PFC connectivity (inter-areal modulation, varying 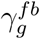, C-D). Each heat map shows the Pearson correlation coefficient 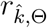 for every combination of the other two parameters. Correlations were calculated over the full range of 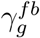 (C) and the largest continuous subset of values showing a reduction in overload (D, see text). White dots in panels A and D show configurations with a significant negative correlation (*p <* 0.05), where peak capacity and overload are consistent with behavioural data from WM tasks, and effective encoding in *PPC* is consistent with neural data from visual tasks (see text, Section 3.2.1).

**Figure 9:**
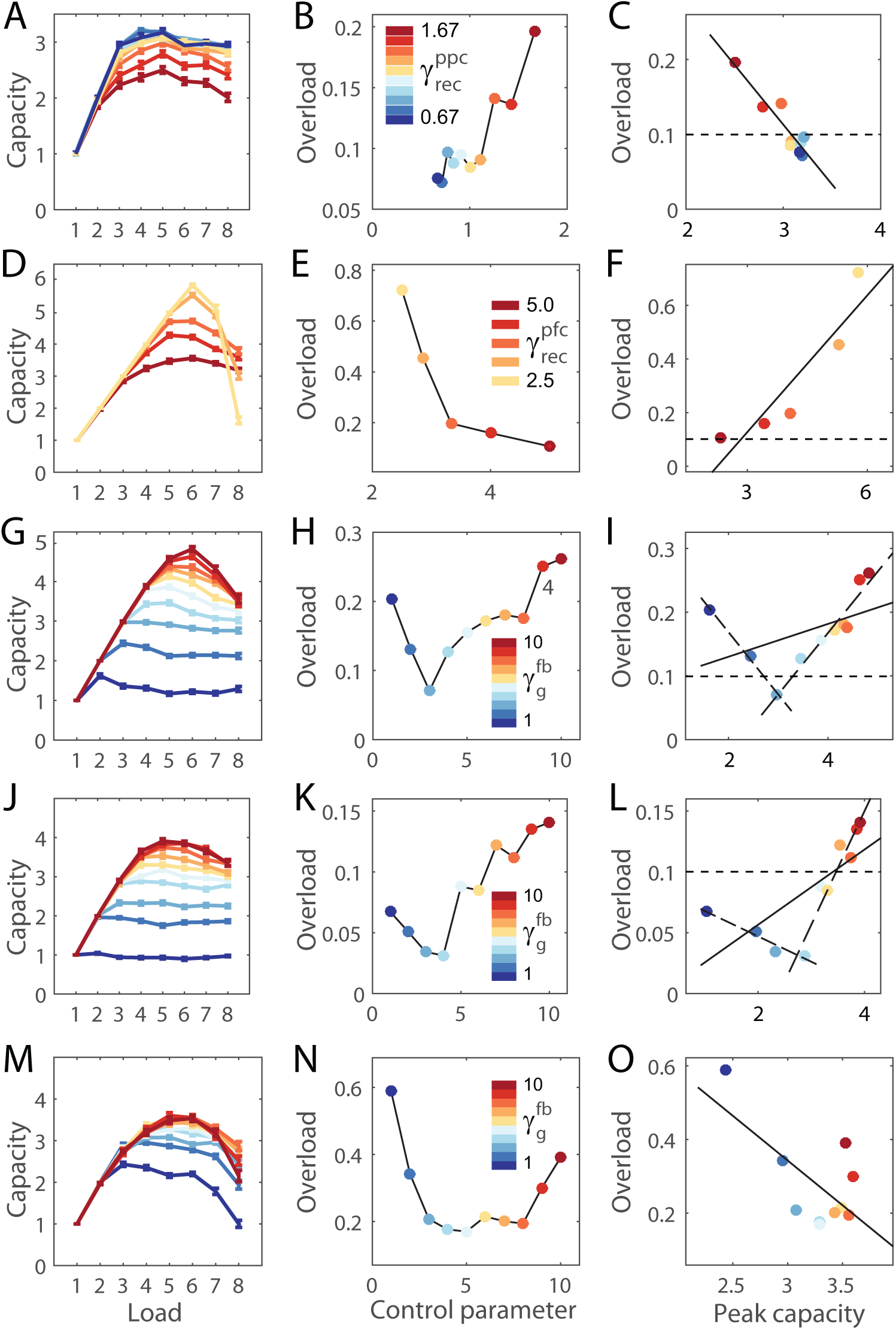
*(preceding page)*: Examples of PPC modulation (first row), PFC modulation (second row) and inter-areal modulation (third to fifth rows) under a single configuration of the remaining two parameters. First, second and third columns show capacity as a function of load, overload as a function of control parameter (given by legend, inset) and overload as a function of peak capacity respectively. (A-C) PPC modulation with 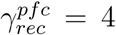 and 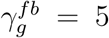. Weaker PPC dynamics increase peak capacity (A) and reduce overload (B), producing a negative correlation between these measures (*r* = −0.935, *p* = 1.000*e* - 4, C). The line in panel C shows the best linear fit (least squares). (D-F) PFC modulation with 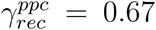 and 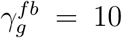. Stronger PFC dynamics reduce overload (E) at the expense of peak capacity (D), producing a positive correlation between these measures (*r* = 0.916, *p* = 0.029, F), as in the PPC-only model. (G-H) Inter-areal modulation with 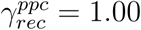 and 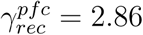. Stronger feedback projections to PPC from PFC increase capacity (G) and decrease overload (H) over a continuous subset of 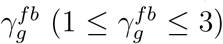, producing a negative correlation (*r* = −0.998, *p* = 0.045, short-dashed line in I). Stronger feedback projections increase capacity and overload, producing a positive correlation over another continuous subset (*r* = 0.961, *p* = 1.000*e* - 4, long-dashed line in I). The solid line in panel I shows the linear fit to the full range of 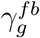. The horizontal dotted line shows Θ = 0.1, chosen as the threshold for alleviating overload (see text). (J-L) Inter-areal modulation with 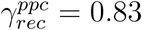 and 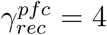. Stronger feedback projections increase capacity (J) and decrease overload (K) over a continuous subset of 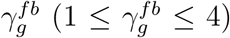, producing a negative correlation (*r* = −0.969, *p* = 0.031, short-dashed line in L) but the magnitude of overload to be overcome is negligible (Θ < 0.07 for all four values fit by the short-dashed line in L). Stronger feedback projections increase capacity and overload over another continuous subset, producing a positive correlation (*r* = 0.948, *p* = 0.001, long-dashed line in L). (M-O) Inter-areal modulation with 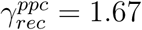 and 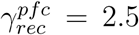. Stronger feedback projections increase capacity (M) and decrease overload (N) over a continuous subset of 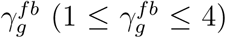, before overload reverses direction with further increases in 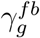. Overload is greatly reduced in magnitude, but never reaches a value consistent with high-capacity subjects and conditions (Θ > 0.17 for all 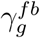).

PFC modulation yielded positive correlations over the full 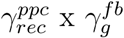 parameter space (Figure 8B). This result was predictable, since limiting the number of items available for storage limits peak capacity. In other words, 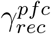 embodies a trade-off between peak capacity and overload, as seen in the PPC-only model (Figure 5). Stronger competitive encoding therefore had a monotonic effect on overload (*e.g.* Figure 9F).

Inter-areal modulation yielded a strong negative correlation (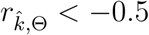 for 6 out of 50 combinations of 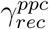 and 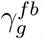) in the corner of the 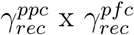 parameter space where stronger recurrent dynamics in *PPC* coincide with weaker recurrent dynamics *PFC* (Figure 8C). Thus, a negative correlation was seen when *PFC* was insufficiently selective and *PPC* was too competitive to be overcome by weak (inter-areal) feedback projections. Stronger feedback projections therefore increased peak capacity and decreased overload by providing more support to the subset of items selected by *PFC*. Notably, increasing the strength of PPC-PFC connectivity rarely had a monotonic effect on overload, so we calculated the correlation between peak capacity and overload for a continuous subset of 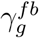, where this subset ranged from 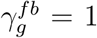 to the value of 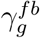 producing the least overload for a given combination of 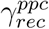 and 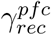 (requiring at least three values, Figure 9I). A strong negative correlation 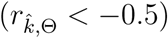 occurred for 18 out of 50 combinations of 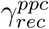 and 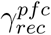, most of which resulted from a broadening of the parameter region identified above (strong and weak recurrent dynamics in *PPC* and *PFC* respectively; compare panels C and D in Figure 8). The rest were spurious, where the magnitude of overload to be overcome was negligible (*e.g.* overload was always below Θ = 0.1 for the negative correlation in Figure 9J-L) or where overload was considerably reduced, but remained high (*e.g.* overload did not drop below Θ = 0.1 in Figure 9M-O).

#### 3.2.2 PPC modulation and inter-aerial modulation alleviate moderate overload and satisfy quantitative constraints

Next, we sought to determine whether peak capacity and the reduction in overload were quantitatively consistent with behavioural data in the parameter regions showing a significant negative correlation (*p <* 0.05), and whether *PPC* was consistent with neural data. As shown in Figure 3, peak capacity should exceed around three items [see Cowan (2001); Luck and Vogel (2013)] and overload should be reduced to close to zero (Chee & Chuah, 2007; Linke et al., 2011; Fukuda, Woodman, &; Vogel, 2015). Additionally, *PPC* should initially encode all stimuli (Thomas & Paré, 2007). For the last of these requirements, we defined *E*^′^ as the minimum ratio of the effective load to the actual load over all load conditions, that is, *E*^′^ = *min*[*E*(*n*)*/n*] (in practice, *E*^′^ = *E*(8)*/*8 under all configurations). Allowing a tolerance of 10%, we therefore searched for parameter regions in which 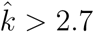, *min*(Θ) < 0.1 and *E*^′^ > 0.9. We included the additional constraint that the magnitude of overload to be overcome [*max*(Θ)] must be greater than 0.1. Under PPC modulation, these criteria were satisfied by five contiguous locations in the parameter region showing a negative correlation (white dots in Figure 8A). Under inter-aerial modulation, they were not satisfied in any parameter region, but they were satisfied in one location of the 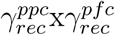 parameter space when we searched the continuous subsets of 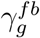 described above (white dot in Figure 8D). These results were unchanged when we lowered our criterion for peak capacity to 2, again with a tolerance of 10% 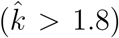. Thus, in isolation, PPC modulation and inter-areal modulation were both able to increase capacity and reduce overload in a manner consistent with neural and behavioural data from multiple-item visual and WM tasks, where weaker PPC dynamics and stronger feedback projections supported better task performance. PPC modulation was the more robust mechanism, as functional connectivity was only able to account for these data under a single combination of the other two parameters.

#### 3.2.3 Control of distributed network dynamics accounts for group differences in studies showing pronounced overload

Finally, we considered our control parameters in combination with one another, aiming to provide a better quantitative account of the magnitude of overload shown by low-capacity subjects, and in low-capacity tasks and conditions (henceforth low-capacity performance). Under PPC modulation and inter-areal modulation, the most extreme cases of overload in the parameter regions satisfying our criteria were Θ ≈ 0.2 (Figure 9A-C and G-I, and Figure 10). As shown in Figure10 (PPC modulation and inter-aerial modulation), this degree of overload is consistent with some data [*e.g.* Cusack et al. (2009); Linke et al. (2011)], but overload has approached 50% of peak capacity in several studies [*e.g.* Chee and Chuah (2007); Xu (2007); Fukuda, Woodman, and Vogel (2015)] and has even approached 100% among elderly, low-capacity subjects in extreme load conditions (Matsuyoshi et al., 2014). We therefore investigated whether simultaneous variation of our control parameters could alleviate the more extreme cases of overload that occur with low-capacity performance, resulting in peak capacity consistent with high-capacity performance in the same studies. High-capacity performance is explained by the computational principles identified above: strong recurrent dynamics in PFC, supporting competitive encoding; strong feedback projections to PPC from PFC, supporting persistent activity; and weak recurrent dynamics in PPC, limiting competition during the memory delay. These principles not only capture a tangible strategy for WM storage on multiple-item tasks (selection of a subset of items for storage), but offer a specific set of mechanisms for their implementation in fronto-parietal circuitry. Thus, we reasoned that low-capacity performance would be explained by *non*-compliance with these principles, investigating the PPC-PFC model under the lowest value of 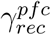, the lowest value of 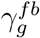 and the highest value of 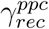 (the low-capacity configuration). In other words, the low-capacity configuration violated the above principles by instantiating the weakest recurrent dynamics in *PFC*, the weakest feedback projections to *PPC* and the strongest recurrent dynamics in *PPC*.

**Figure 10:**
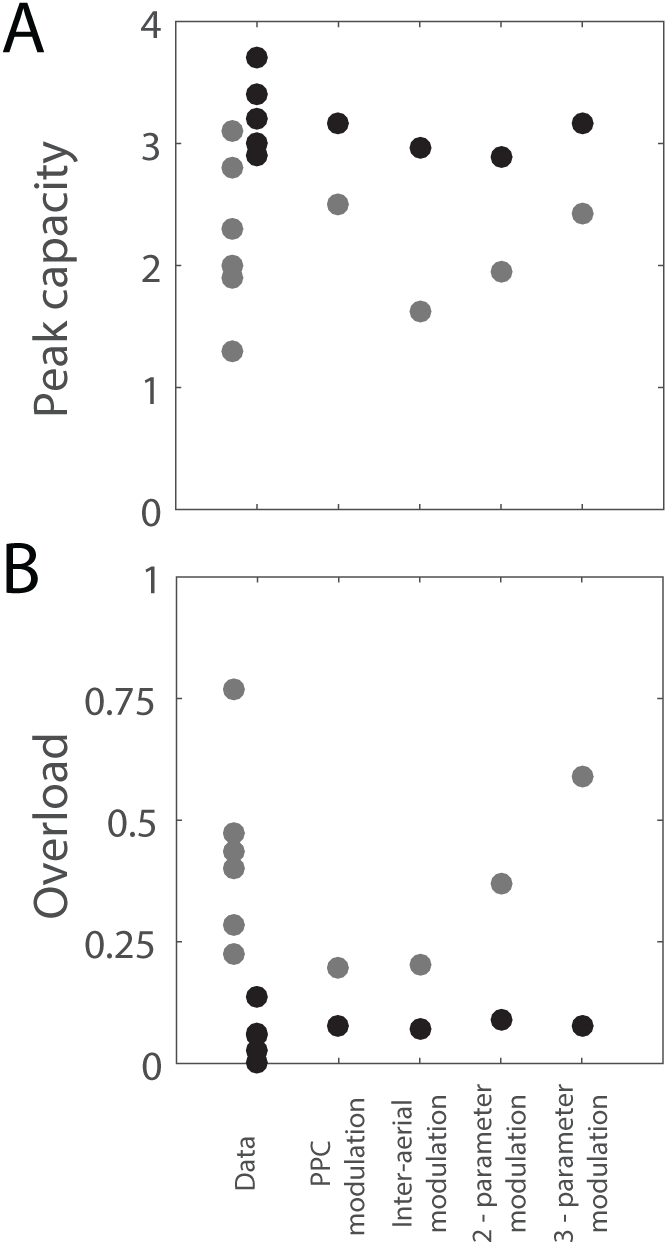
Peak capacity (A) and overload (B) corresponding to lowcapacity (grey) and high-capacity (black) performance by experimental subjects (Data, left) and by the PPC-PFC model under PPC modulation (middle left), inter-aerial modulation (middle), 2-parameter modulation (middle right), and 3-parameter modulation (right). Experimental data are reproduced from Figure 3. Low-capacity and high-capacity data are horizontally staggered for clarity. Low and high-capacity configurations under PPC modulation correspond to 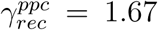 (strongest) and 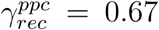 (weakest), shown in Figure 11A-C. Low and high-capacity configurations under interaerial modulation correspond to 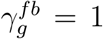 (weakest) and 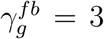, shown in Figure 11G-I. Low and high-capacity configurations under 2-parameter and 3-parameter modulation correspond to Figures 11D-F and A-C respectively.

The low-capacity configuration had a peak capacity of 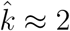 items, overload of Θ > 0.5 and allowed all items to be encoded by *PPC* (Figure 11A-C, grey). For a range of parameter values, increasing the strength of competition in *PFC* (increasing 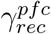), increasing the strength of feedback projections to *PPC* (increasing 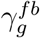) and decreasing the strength of competition in *PPC* (decreasing 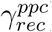) produced very similar results to those shown in Figure 7 (high-capacity configurations), raising peak capacity to 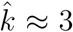 items, reducing overload to close to Θ = 0 and allowing the encoding of all items in *PPC*. These results are strikingly similar to peak capacity and overload among low- and high-capacity subjects [*cf.* Fukuda, Woodman, and Vogel (2015)] and task conditions [*cf.* Chee and Chuah (2007)] (Figure 10). We do not propose a specific trajectory through the parameter space from one extreme to the other, but rather, we emphasize that multiple trajectories show robustness of the computational principles identified by our simulations.

**Figure 11:**
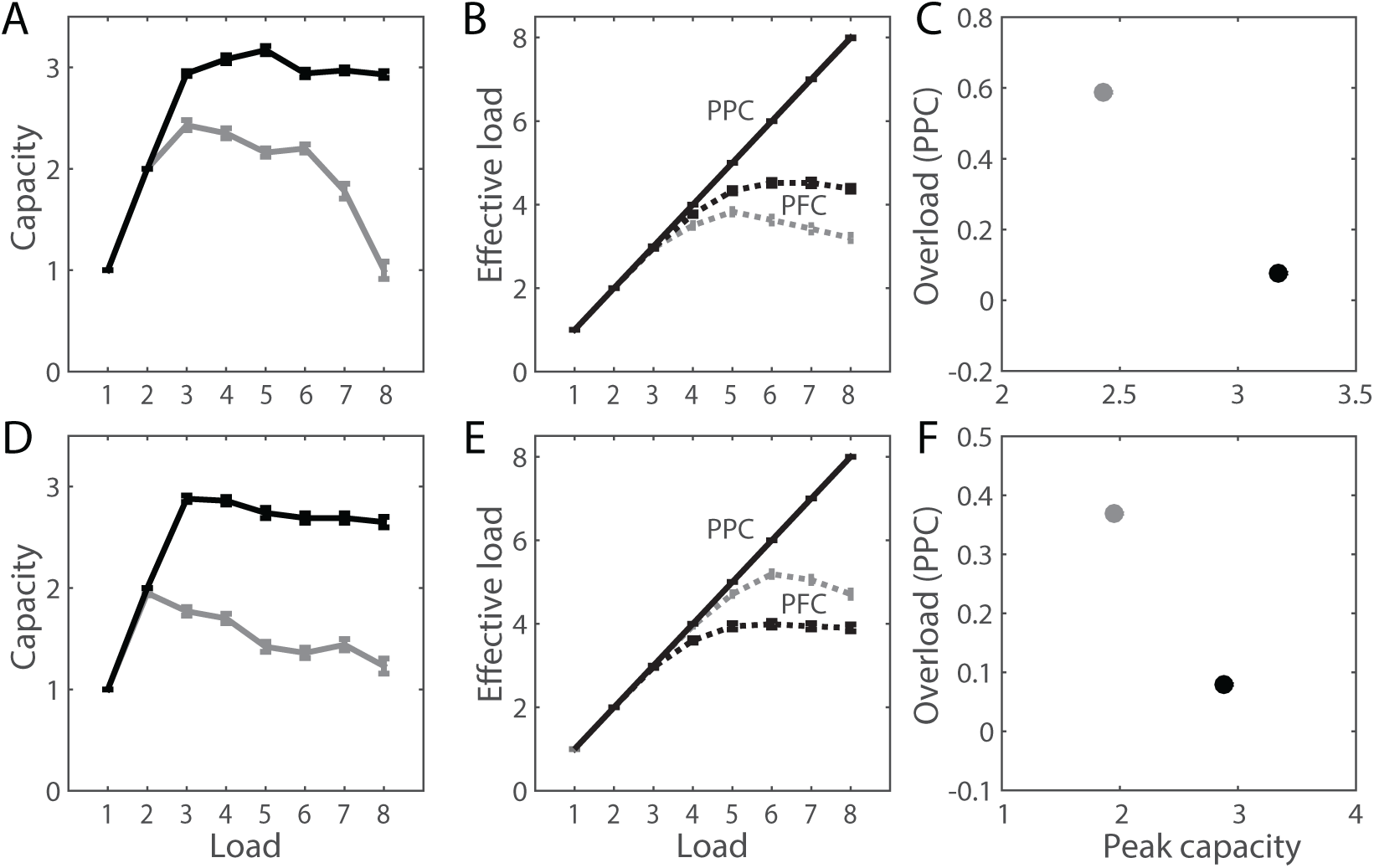
(A-C) Simultaneously increasing the strength of competition in *PFC*, increasing the strength of feedback projections to *PPC* from *PFC*, and decreasing the strength of competition in *PPC* (3-parameter modulation) raised peak capacity (A) and reduced overload (C) in a manner consistent with neural and behavioural data (see text, Section 3.2.1). Effective load in *PFC* (dotted curves in panel B) was higher under the high-capacity configuration (black, parameter values provided in Figure 7) than the low-capacity configuration (grey; 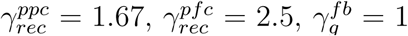). 3-parameter modulation was therefore inconsistent with the hypothesis that low capacity and pronounced overload result from the encoding of too many items in *PFC*. (D-F) Simultaneously increasing the strength of competition in *PFC*, increasing the strength of feedback projections to *PPC*, and fixing *PPC* dynamics at a moderate level (2-parameter modulation) similarly raised peak capacity and reduced overload, but the effective load in *PFC* was higher under the low-capacity (grey; 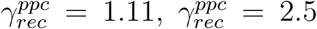 and 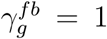) configuration than under the high-capacity (black; 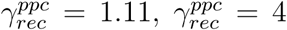 and 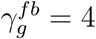) configuration (E). 2-parameter modulation was therefore consistent with the hypothesis that low capacity and pronounced overload result from the encoding of too many items in *PFC*.

Surprisingly, under the high-capacity configurations satisfying the criteria described in Section 3.2.2 (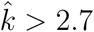, *min*(Θ) < 0.1 and *E*^′^ > 0.9), modulation of all three parameters (henceforth 3-parameter modulation) produced an effective load in *PFC* that was greater than or equal to the effective load under the low-parameter configuration (compare the black and grey dashed curves in Figure 11B). This result does not support the hypothesis that poor performance by low-capacity subjects results from a failure to select a manageable subset of items for storage (poor task strategy) since stimulus encoding in *PFC* was at least as selective under the low-capacity configuration as under the high-capacity configurations. Rather, it supports the hypothesis that low-capacity and pronounced overload reflect poor control over fronto-parietal circuitry, *i.e.* a relative inability to strengthen competitive dynamics in PFC, weaken dynamics in PPC, and enhance the functional coupling between these regions. We therefore determined whether the simultaneous modulation of any two parameters (2-parameter modulation, holding the third parameter fixed) could satisfy the above criteria, overcome a degree of overload comparable to that of the low-capacity configuration (Θ ≈ 0.5), and show a *lower* effective load in *PFC* with high capacity than with the low capacity. If so, such a configuration would support the hypothesis that low-capacity subjects are encoding too many memoranda and would allow us to consider predictions under 3-parameter and 2-parameter modulation that might distinguish between these two hypotheses (poor task strategy *vs.* poor cognitive control). To this end, we simultaneously decreased and increased 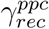 and 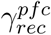 from their highest and lowest values respectively, while holding 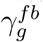 fixed (at all possible values); simultaneously decreased and increased 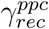 and 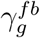 from their highest and lowest values respectively, while holding 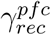 fixed; and simultaneously increased 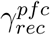 and 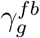 from their lowest values, while holding 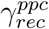 fixed. The first two approaches were unable to satisfy our constraints, but the third approach was able to do so with a moderate increase in the strength of competition in *PFC*, a small increase in the strength of feedback projections, and with *PPC* dynamics fixed at a moderate level (Figure 11D-F).

#### 3.2.4 Simulated EEG recordings resolve conflicting hypotheses on working memory performance captured by the PPC-PFC model

Having determined that 3-parameter and 2-parameter modulation offer competing explanations for the behavioural phenomenon of WM overload, we sought to distinguish between these explanations by determining their respective abilities to account for neural data from an experimental task showing overload. To the best of our knowledge, the only such data currently available are the EEG recordings over parieto-occipital cortex by Fukuda, Woodman, and Vogel (2015). Thus, we approximated EEG recordings over PPC and lateral PFC under 3-parameter and 2-parameter modulation. To approximate the EEG signal, we followed the approach by McCarthy, Brown, and Kopell (2008), summing all excitatory currents onto pyramidal neurons in each network. Thus, we simulated the instantaneous source amplitude of EEG over PPC by 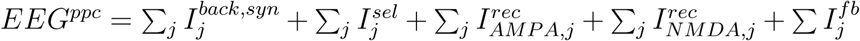 and over lateral PFC by 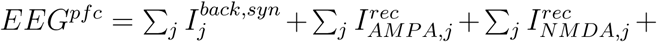 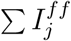, where index *j* refers to pyramidal neurons, currents 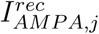 and 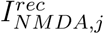 are given by Equation 6, and currents 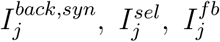 and 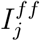 are given by Equations 9, 18, 16 and 15 respectively. We summed this instantaneous signal over the portion of the memory delay used to determine WM performance (the last 300ms, Section 2.5) to obtain the total amplitude 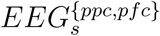 during this time window. Because competitive dynamics in PPC and PFC may be modulated by different mechanisms than the scaling of synaptic conductances used here (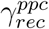 and 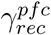) and because our EEG approximation sums the resulting synaptic currents, we normalized the EEG approximation by its minimum and maximum values (*EEG*_*η*_ = [*EEG*_*s*_ - *min*(*EEG*_*s*_)]*/*[*max*(*EEG*_*s*_) - *min*(*EEG*_*s*_)]), predicting the qualitative form of EEG amplitude as a function of memory load, rather than absolute amplitude.

*EEG*_*η*_ was qualitatively distinct during the delay interval under 3-parameter and 2-parameter modulation. Under 3-parameter modulation, *EEG*_*η*_ was bi-linear over memory load for both networks under low- and high-capacity configurations, where the ‘second line’ of the bi-linear curve had a negative slope under the low-capacity configuration and was approximately horizontal under the high-capacity configuration (Figure 12A). These curves are strikingly similar to the EEG amplitude shown by Fukuda, Woodman, and Vogel (2015) for low- and high-capacity subjects (their Figure 4C). Under 2-parameter modulation, *EEG*_*η*_ was tri-linear for both networks (Figure 12B) and therefore did not account for the available EEG data. As such, these findings are strongly supportive of the principles of 3-parameter modulation and its corresponding hypothesis that low-capacity subjects are indeed selecting a manageable subset of items for storage, but that they have poor control over their fronto-parietal circuitry (corresponding to 3-parameter modulation here).

**Figure 12:**
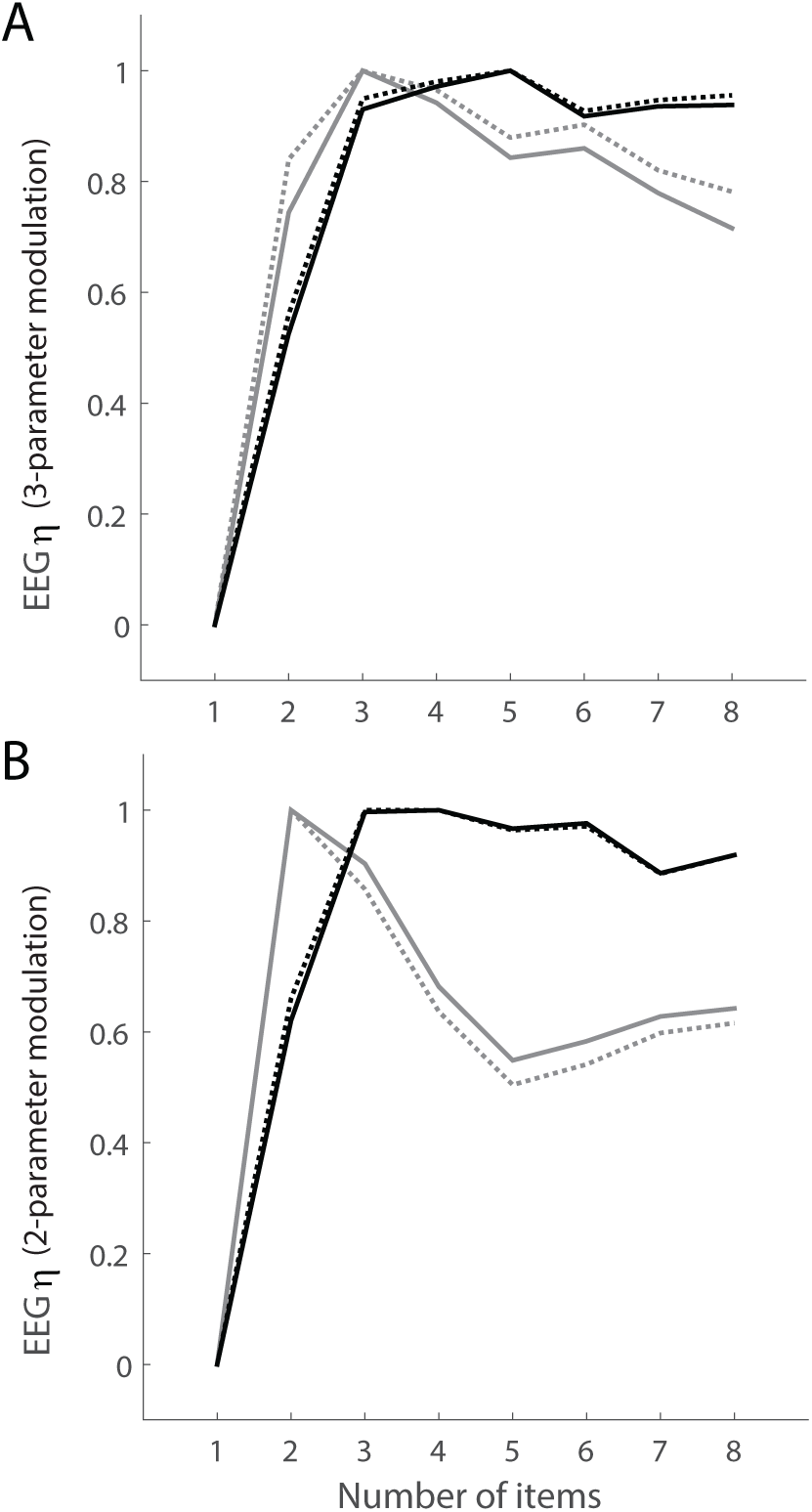
Normalized EEG approximation (*EEG*_*η*_, see text) for the PPC (solid) and PFC (dotted) networks under the low-capacity (grey) and high-capacity (black) configurations over the last 300ms of the memory delay (used to determine WM performance in the model). (A) Under 3-parameter modulation, *EEG*_*η*_ was bilinear for both networks under the low- and high-capacity conditions, where the slope of the ‘second line’ was negative under the low-capacity configuration [*cf.* Fukuda et al (2015)]. (B) Under 2-parameter modulation (*PPC* dynamics fixed at a moderate level, see text), *EEG*_*η*_ for both networks was again bi-linear under the high-capacity configuration, but was tri-linear under the low-capacity configuration.

To test the robustness of these results and to facilitate a more direct comparison with the EEG data by Fukuda, Woodman, and Vogel (2015), we calculated *EEG*_*η*_ over longer windows at the end of the memory delay. These authors used a 150ms stimulus interval and a 1s delay interval on their task, averaging the signals from parieto-occipital channels over the last 850ms of the delay. We therefore calculated *EEG*_*η*_ over a series of increasingly long time windows (steps of 50ms), ranging from the last 300ms to the last 850ms of the memory delay. Our results were qualitatively robust for time windows of up to ∼750ms (not shown).

#### 3.2.5 An EEG signature for hierarchical recruitment of competition during stimulus encoding

Having shown that hierarchical recruitment of competition during stimulus encoding accounts for behavioural (Figures 3 and 11) and neural (Figure 12) data from multiple-item WM tasks on which memory load exceeds subjects’ retention abilities, we sought to identify a measurable signature of this hypothesis. We therefore calculated *EEG*_*η*_ during the stimulus interval under 3-parameter modulation, having ruled out 2-parameter modulation from further consideration, due to its inconsistency with delay-interval EEG activity (previous section). During the stimulus interval, *EEG*_*η*_ showed greater concavity (concave down) as a function of memory load for *PFC* than *PPC* (Figure 13) under the low- and high-capacity configurations alike. In this regard, concavity serves an index of the timing of selective encoding, *i.e.* greater concavity reveals earlier selection of memoranda. Thus the PPC-PFC model makes a specific, testable prediction for our hypothesis: EEG amplitude will show greater concavity over memory load when recorded over lateral PFC than when recorded over PPC during the stimulus interval of multiple-item WM tasks.

**Figure 13:**
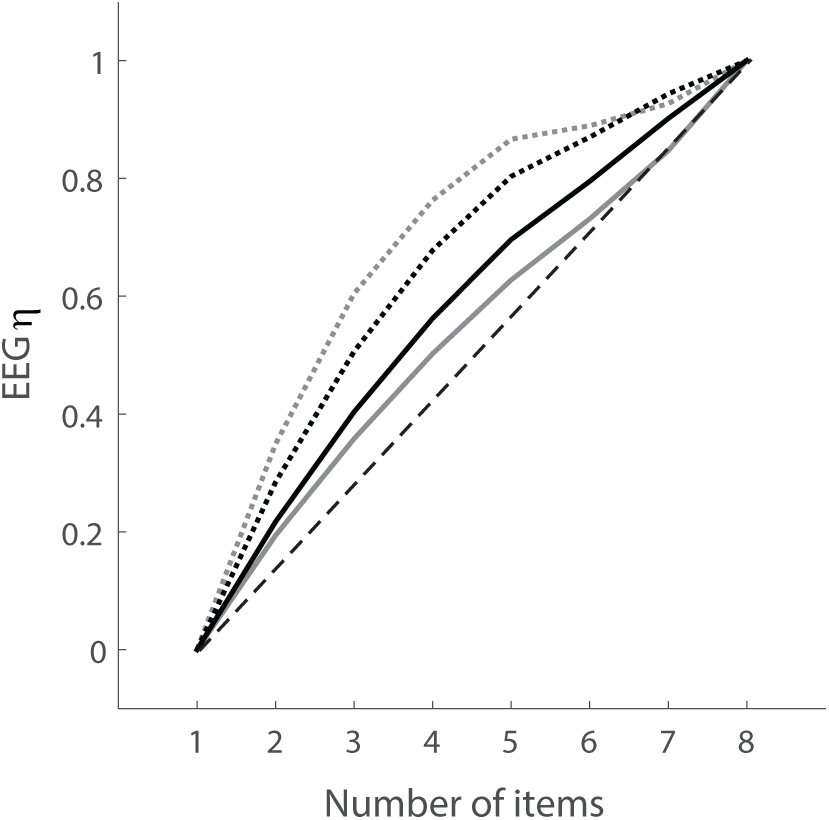
Predictions by the PPC-PFC model for normalized EEG amplitude (*EEG*_*η*_, see text) over PPC (solid) and lateral PFC (dotted) for low-capacity (grey) and high-capacity (black) WM performance during the stimulus interval. Under 3-parameter modulation, hierarchical recruitment of competition during stimulus encoding predicts greater concavity over PFC, indicating earlier selection of memoranda than in PPC. The dashed unity line highlights concavity of the curves.

## 4 Discussion

The storage limitations of WM have been the subject of intense research interest for several decades [see Luck and Vogel (2013)], but although several studies have reported a reduction in WM capacity with high memory load [*e.g.* Xu (2007); Chee and Chuah (2007)], WM overload has only been the focus of a handful of behavioural experiments (Cusack et al., 2009; Linke et al., 2011; Matsuyoshi et al., 2014; Fukuda, Woodman, & Vogel, 2015). We investigated the neural basis of overload with the PPC and PPC-PFC models, finding that overload could be reduced in both models by strong competitive dynamics during the stimulus interval of simulated WM tasks. The PPC-only model, however, showed a positive correlation between peak capacity and overload (Figure 5), in opposition to available data (Figure 3).

The PPC-PFC model accounted for these data in a parameter regime where selective encoding was supported by strong competitive dynamics in *PFC*, persistent activity was supported by inter-areal projections, and weak dynamics in *PPC* limited competition during the memory delay (Figure 7). As such, the model implemented hierarchical recruitment of competition during stimulus encoding and identified a set of computational principles for WM storage in distributed circuitry. Under these principles, all WM items were encoded by *PPC* (Figure 7B), consistent with single-cell electrophysiological recordings from PPC (Thomas & Paré, 2007); simulated EEG amplitude was bi-linear over memory load during the delay period (Figure 12A, black curves), consistent with EEG recordings over parieto-occipital cortex (Vogel & Machizawa, 2004; Fukuda, Woodman, & Vogel, 2015); and peak capacity was around three items, consistent with behavioural data from numerous WM tasks (Figure 7B) [see Cowan (2001); Luck and Vogel (2013)]. When we violated the identified principles (increased competition in *PPC*, decreased strength of feedback projections and decreased competition in *PFC*), peak capacity was reduced to just over two items (Figure 11A), overload was greater than 50% of peak capacity (Figure 11C) and the ‘second line’ of the bi-linearity of simulated EEG amplitude showed a negative slope (Figure 12A, grey curves). These results are strikingly consistent with behavioural and EEG data from low-capacity subjects in the study by Fukuda, Woodman, and Vogel (2015). To our surprise, the model implemented selective encoding in this low-capacity regime (Figure 11B). Thus, while it captured a strategy for WM storage under high load and offered a set of neural mechanisms for its implementation in hierarchical circuitry, it predicted that low-capacity subjects are indeed attempting this strategy and that their performance reflects poor control of fronto-parietal processing. Our hypothesis is testable by the prediction that EEG amplitude over memory load will show greater concavity over lateral PFC than over PPC during the stimulus interval of WM tasks (Figure 13), providing a neural signature of early selection.

### 4.1 Limitations of our models

Our models have limitations, of course. While there is growing support for the hypotheses that PPC is the hub of distributed WM storage (Palva et al., 2010; Christophel et al., 2012; Salazar et al., 2012) and that fronto-parietal interactions play a central role in multiple-item storage (Edin et al., 2009; Palva et al., 2010), other brain regions play important roles in WM [see Sreenivasan, Curtis, and D’Esposito (2014); D’Esposito and Postle (2015)]. Thus, we do not claim that the PPC-PFC model and the computational principles it identifies should explain WM overload under all possible conditions. For example, the negative correlation between peak capacity and overload that guided our investigations (Figure 3) is common (Chee & Chuah, 2007; Xu, 2007; Cusack et al., 2009; Linke et al., 2011; Matsuyoshi et al., 2014; Fukuda, Woodman, & Vogel, 2015), but not ubiquitous. These two measures have been shown to increase together with the duration of stimulus encoding [see Figure 4 by Cusack et al. (2009)], as well as during childhood development [from 6–7 year-old children to college students in the study by Cowan, Morey, AuBuchon, Zwilling, and Gilchrist (2010)].

Another limitation of our models is that they only consider the spatial location of memoranda, ignoring other features and their conjunctions. In effect, our simulations assume that everything encoded by PPC satisfies a set of rules for selection, *e.g.* red squares or blue circles. This approach is common among neural models of WM storage [*e.g.* Compte et al. (2000); Tanaka (2002); Macoveanu, Klingberg, and Tegnér (2006); Edin et al. (2009); Wei et al. (2012); Standage and Paré (2018)] and is reasonable for studies addressing capacity. While the PPC-PFC model takes an important step toward the understanding of WM storage in distributed circuitry, an understanding of feature-bound memoranda will likely require hierarchical models with converging feature maps [*e.g.* Swan and Wyble (2014)]. See Raffone and Wolters (2001) for a binding mechanism for sequentially presented memoranda [related models are described by Lisman and Idiart (1995); Jensen and Lisman (1996)]. Additionally, the target stimuli in our simulations were equidistant from one another, which was not the case under most conditions of the experiments providing the data to which we compare model performance (Chee & Chuah, 2007; Xu, 2007; Cusack et al., 2009; Linke et al., 2011; Matsuyoshi et al., 2014; Fukuda, Woodman, & Vogel, 2015). We used equidistant targets in order to keep our simulations as simple as possible (and our results as interpretable as possible), bearing in mind that the PPC-PFC model is comprised of bidirectionally-coupled dynamic systems. To the best of our knowledge, ours is the first study to systematically investigate the neural basis of WM overload and we have focused on its generalities across different experimental approaches. Future studies should investigate overload under the specific conditions of individual experiments, including the spatial clustering of stimuli. For commentary of the effects of spatial clustering on WM performance, see Standage and Paré (2018).

Finally, our focus on WM overload inherently limits our study to the investigation of capacity, but we do not suggest that capacity provides the only limitation on WM storage. There is ample evidence that the precision of memoranda is load-dependent [*e.g.* Zhang and Luck (2008); Bays, Catalao, and Husain (2009); van den Berg, Shin, Chou, George, and Ma (2012); Schneegans and Bays (2016)]. Historically, capacity and precision have been presented as evidence for conflicting hypotheses on the nature of WM storage [see Luck and Vogel (2013); Ma et al. (2014)], but neural modelling studies have begun to focus on their relationship and its neural basis (Wei et al., 2012; Roggeman, Klingberg, Feenstra, Compte, & Almeida, 2013; Okimura, Tanaka, Maeda, Kato, & Mimura, 2015; Standage & Paré, 2018). We are unaware of studies showing anything resembling overload in relation to precision (*e.g.* unchanging precision up to a critical load, followed by a decrease), but future experiments should investigate the dependence of precision on supra-capacity memory load.

### 4.2 Persistent activity and distributed working memory storage

While our simulations support the hypothesis that inter-aerial projections between PPC and PFC support persistent mnemonic activity, we do not suggest that persistent activity is the only mechanism by which target stimuli may be stored over a delay interval, nor that target stimuli provide the only task-relevant information stored by persistent activity. There is a growing body of evidence for ‘state-based’ hypotheses of WM storage, which posit that the same neural populations represent WM targets before and after stimulus offset, and that attention determines their state of activation after offset. This general principle is supported by the ‘synaptic theory of working memory’, according to which, a non-selective signal refreshes synaptic traces among stimulus-encoding neural populations, reactivating these populations following a memory delay (Mongillo, Barak, & Tsodyks, 2008). In this model, the non-selective signal plays the role of attention and short term synaptic facilitation allows synaptic traces to persist for around 1s. Whether the timescales of short term facilitation and depression in sensory cortices are compatible with this mechanism is debatable, but the model provides a compelling proof of concept for the implementation of state-based WM storage. There is also a growing body of evidence for the encoding of various kinds of task-relevant information by persistent activity, such as task rules and the categories of memoranda [see Sreenivasan et al. (2014)]. It seems unlikely that any one mechanism should account for all aspects of a construct as broad and nuanced as WM, and our hypothesis that hierarchical recruitment of competition during stimulus encoding ameliorates overload is by no means a hypothesis against other mechanistic explanations of other data. Quite the opposite, it offers a compatible selection mechanism to more traditional notions of top-down attentional signals to lower cortices [see D’Esposito and Postle (2015)].

Along a similar vein, the computational principles of our hierarchical model are not necessarily limited to PPC and PFC, and may be applicable to any bidirectionally-coupled regions in the cortical hierarchy. While our criteria for the selection of parameter configurations in Section 3.2.1 included the encoding of all target stimuli by *PPC* (with a tolerance of 10%) and therefore favoured weaker, less competitive dynamics in *PPC* than *PFC*, it is plausible that recurrent dynamics increase in strength with hierarchical ascendancy more generally. This possibility is consistent with systematic variation in pyramidal cell morphology with hierarchical ascendancy, such as increased spine density and dendritic branching (Elston, 2002). In this regard, our prediction that EEG amplitude over PPC during stimulus encoding will show greater concavity as a function of memory load than over PFC may serve as a more general signature for hierarchical recruitment of competition. Nonetheless, it is important to reiterate that neural data from PPC provided a strong constraint on the parameter values of the PPC-PFC model and that the extensive body of data pointing to fronto-parietal involvement in WM storage [see Eriksson, Vogel, Lansner, Bergstrom, and Nyberg (2003); Constantinidis and Klingberg (2016); Christophel, Klink, Spitzer, Roelfsema, and Haynes (2017)] provides good reason to ground the model in fronto-parietal areas. It is also worth noting that spike rates in *PFC* were much lower lower than in *PPC* in the model (Figure 6), consistent with the relative spike rates of these cortical areas *in vivo* [*e.g.* Swaminathan and Freedman (2012); Murray et al. (2014)]. We do not claim that PFC spike rates are low because inhibition is stronger in PFC than in other cortical areas (*e.g.* PPC) but nonetheless, we emphasize that the PPC-PFC model is not only consistent with behavioural (Figure 10) and EEG (12) data from studies showing WM overload, but also with single-cell recordings from PPC [*e.g.* Funahashi, Bruce, and Goldman-Rakic (1989); Takeda and Funahashi (2002); M. Wang et al. (2011)] and PFC [*e.g.* Gnadt and Andersen (1988); Paré and Wurtz (1997)] in non-human primate studies.

### 4.3 Beyond local-circuit attractor models

Local-circuit attractor models (such as the PPC-only model) have been invaluable to our understanding of the neural basis of persistent activity on single-item tasks (X.-J. Wang, 1999; Compte et al., 2000) and capacity limitations on multiple-item tasks with memory loads similar to (or less than) capacity (Tanaka, 2002; Macoveanu et al., 2006; Edin et al., 2009). Assuming that all items are encoded for storage, these models necessarily produce overload when the number of memoranda sufficiently exceeds capacity, due to the competition between simulated neural populations [see Edin et al. (2009) for analysis]. This finding reveals a limitation of local-circuit models of WM storage, since not all experimental tasks, conditions and subjects show overload [*e.g.* Xu (2007); Chee and Chuah (2007); Cusack et al. (2009); Fukuda, Woodman, and Vogel (2015)]. The same can be said of hierarchical models in which a top-down control signal modulates the recurrent dynamics of a downstream network (Edin et al., 2009; Roggeman et al., 2013), since persistent activity is supported by attractor dynamics in the network receiving the control signal (Edin et al., 2009). The PPC-PFC model builds on this work, taking a step toward an understanding of the roles played by local circuits in distributed WM storage. An important next step is to simulate inter-areal cortical pathways in more detail, since these pathways systematically differ according to layer, hierarchical distance and (presumably) function. The structural and mechanistic differences between the PPC-PFC model and the model by Edin et al. (2009) are instructive in this regard. Our model emphasizes the role of topographic inter-areal pathways, which run bidirectionally in supra-granular layers between hierarchically adjacent cortical areas (such as PPC and PFC), but which are increasingly dominated by feed-forward (ascending) projections with greater hierarchical distance. In the model by (Edin et al., 2009) [and Roggeman et al. (2013)], the top-down control signal is spatially non-selective (diffuse), an established form of gain modulation in local-circuit models of this class (Salinas & Abbott, 1996; Furman & Wang, 2008; Standage, You, Wang, & Dorris, 2013). Diffuse pathways run bidirectionally in infra-granular layers between adjacent cortical areas, but are increasingly dominated by feedback (descending) projections with greater hierarchical distance, *i.e.* the opposite arrangement to topographic pathways [see Markov and Kennedy (2013)]. Thus, our different approaches capture fundamentally different mechanisms for control of WM storage: bottom-up recruitment by (and of) topographic pathways and top-down control by diffuse pathways respectively. It seems likely that both mechanisms are involved in WM storage. Future work should test the predictions of our respective models, aiming to identify the roles of different cortical areas and their functional interactions in support of WM.

Our EEG approximations with the PPC-PFC model also point to an exciting direction for future research. In the present work, the summation of excitatory currents onto simulated pyramidal neurons (McCarthy et al., 2008) allowed us to approximate the instantaneous EEG source amplitude over PPC and PFC, but we did not use this methodology to investigate the possible role of oscillations in WM storage. In attractor models with synaptic resolution, oscillations can be controlled by the ratio of the time constants of excitatory and inhibitory synaptic receptors (Brunel & Wang, 2003) and therefore by the relative strengths of AMPARs and NMDARs [*e.g.* Compte et al. (2000); Buehlmann and Deco (2008)], since the time constants of these receptors are shorter and longer respectively than the time constant of GABARs (Table 1). In the interest of simplicity, we purposefully avoided oscillations in the PPC-PFC model, but bidirectionally-coupled attractor networks have been used to investigate inter-aerial transmission of information in earlier studies [*e.g.* Buehlmann and Deco (2010)], suggesting that our hierarchical model and simulation paradigm are ideal for investigating the mechanisms by which synchronized (Palva et al., 2010) and desynchronized (Fukuda, Mance, & Vogel, 2015; Fukuda, Kang, & Woodman, 2016) oscillations may contribute to WM storage, and the relationships between these EEG data and oscillations in local field potentials during WM tasks (Lundqvist et al., 2016; Lundqvist, Herman, Warden, Brincat, & Miller, 2018).

## 5 Conclusions

The real-world phenomenon of WM overload has long been of concern to educators (Sweller, 1988), who have identified the need for a stronger scientific foundation for pedagogic strategies aiming to prevent its occurrence in the classroom (Schnotz & Kurschner, 2007; de Jong, 2010). Nonetheless, despite intense research interest in the storage limitations of WM more generally [see Luck and Vogel (2013)], only a handful of studies have specifically investigated overload (Cusack et al., 2009; Linke et al., 2011; Matsuyoshi et al., 2014; Fukuda, Woodman, & Vogel, 2015) and to the best of our knowledge, no previous study has investigated its mechanistic basis. Our findings point to cognitive control as the source of differential WM performance across subject groups, rather than capacity *per se*. This finding is consistent with recent experimental work emphasizing strategic ability as the source of high performance on WM tasks and on tests of cognitive ability more generally (Cusack et al., 2009; Linke et al., 2011). Given the strong correlation between capacity and scores on intelligence tests [see Unsworth et al. (2014)], we believe this message is a positive one, though our findings do not suggest that capacity can necessarily be improved by simple strategic adjustments (Section 3.2.3). Rather, they suggest that individuals with better control of distributed cortical processing are better positioned to implement effective strategies. Significant research investment will be required to identify ways to improve this control. Our predictions for experimental testing (Section 3.2.5) are a step in this direction.

## Footnotes

1 The effective load for the strongest recurrent dynamics in the PPC-only model (highest value of 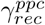, darkest red curves in Figure 5) is less than capacity in some load conditions, i.e. *E*(*n*) *< K*(*n*) (Figure 5A and B). This seeming anomaly reflects the later onset of stimulus-selective activity with such strong recurrent dynamics, where the rate of item-encoding activity (calculated over the full stimulus interval) is too low to satisfy our criteria for encoded items. This effect can be seen in Figure 2, where the onset of item-encoding activity occurs later with higher 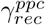.

